# The genomic landscape of contemporary western Remote Oceanians

**DOI:** 10.1101/2022.01.10.475623

**Authors:** Lara R. Arauna, Jacob Bergstedt, Jeremy Choin, Javier Mendoza-Revilla, Christine Harmant, Maguelonne Roux, Alex Mas-Sandoval, Laure Lémée, Heidi Colleran, Alexandre François, Frédérique Valentin, Olivier Cassar, Antoine Gessain, Lluis Quintana-Murci, Etienne Patin

## Abstract

The Vanuatu archipelago served as a gateway to Remote Oceania during one of the most extensive human migrations to uninhabited lands, ~3,000 years ago. Ancient DNA studies suggest an initial settlement by East Asian-related peoples that was quickly followed by the arrival of Papuan-related populations, leading to a major population turnover. Yet, there is uncertainty over the population processes and the sociocultural factors that have shaped the genomic diversity of ni-Vanuatu, who present nowadays among the world’s highest linguistic and cultural diversity. Here, we report new genome-wide data for 1,433 contemporary ni-Vanuatu from 29 different islands, including 287 couples. We find that ni-Vanuatu derive their East Asian- and Papuan-related ancestry from the same source populations and descend from relatively synchronous, sex-biased admixture events that occurred ~1,700-2,300 years ago, indicating a peopling history common to all the archipelago. However, East Asian-related ancestry proportions differ markedly across islands, suggesting that the Papuan-related population turnover was geographically uneven. Furthermore, we detect Polynesian ancestry arriving ~600-1,000 years ago to South Vanuatu in both Polynesian- and non-Polynesian-speaking populations. Lastly, we provide evidence for a tendency of spouses to carry similar genetic ancestry, when accounting for relatedness avoidance. The signal is not driven by strong genetic effects of specific loci or trait-associated variants, suggesting that it results instead from social assortative mating. Altogether, our findings provide insight into both the genetic history of ni-Vanuatu populations and how sociocultural processes have shaped the diversity of their genomes.

## INTRODUCTION

Vanuatu, an archipelago located in western Remote Oceania, is a key region for understanding the peopling history of the Pacific. The cultural, anthropological and genetic diversity of Vanuatu reflects three distinct phases of population movements. The first, which started in present-day Taiwan ~5,000 ya, was associated with the spread of Austronesian languages to Near and Remote Oceania [1–3]. The so-called Austronesian expansion led to the emergence of the well-characterized Lapita cultural complex, which emerged in the Bismarck Archipelago and reached Vanuatu ~3,000 ya [4–6]. Morphometric and ancient DNA (aDNA) studies indicate that the Lapita people in Vanuatu carried East Asian-related ancestry, supporting a connection with the Austronesian expansion [7, 8]. The second migration occurred after the Lapita period, ~2,500 ya, and involved the arrival of Papuan-related peoples who shared ancestry with contemporary Bismarck Archipelago islanders. aDNA studies have shown that these migrations triggered a dramatic shift in genetic ancestry, from the East Asian-related ancestry observed in first Remote Oceanians to the Papuan-related ancestry that has remained predominant since then [9, 10]. Finally, it has been postulated that “Polynesian outliers” from Vanuatu (i.e., Polynesian speakers living outside the Polynesia triangle) are the descendants of migrations from Polynesia into western Remote Oceania [11, 12], a model that has received recent genetic support [13].

Although aDNA studies have revealed that ni-Vanuatu are descended from at least three ancestral populations [9, 10, 13], whether the settlement process was uniform across the multiple islands of the archipelago remains an open question. aDNA data indicate that individuals dated to ~2,500 ya from Central and South Vanuatu carried largely different proportions of East Asian-related ancestry [9, 10, 13], in line with either a unique population turnover that was geographically heterogeneous, or separate admixture events between different populations across islands. Furthermore, Vanuatu is the country with the world’s largest number of languages per capita [14], which supports the view that languages have rapidly diversified since the initial settlement and/or that the arrival of new groups to the archipelago further increased linguistic diversity. Nevertheless, these questions have been difficult to resolve because, to date, available ancient and modern DNA data from the region have remained sparse [9, 10].

Here, we generated genome-wide genotype data for 1,433 contemporary ni-Vanuatu and assessed their fine-scale genetic structure, in order to address central questions about the settlement of western Remote Oceania: do all contemporary ni-Vanuatu derive their ancestry from three populations only? Was the contribution of these three ancestral populations different across the archipelago? Was the Post-Lapita shift to Papuan-related ancestry heterogeneous across islands? In addition, the recent settlement of Vanuatu allows for the study of how socio-cultural practices have shaped genetic diversity in humans over the past ~3,000 years, in the form of sex-biased admixture, relatedness avoidance and non-random mating [15–17]. We thus used the comprehensive set of population genomics data presented here, which includes 287 male-female couples, to answer important questions relating to the genetic history of a population that descends from diverse ancestral populations: Was admixture sex-biased? Was admixture accompanied by language shifts? Has socio-cultural structure influenced mating? Can residence rules and urbanization affect human genetic structure?

## RESULTS

### The Genetic Variation of Ni-Vanuatu is Spatially Structured

To shed light on the genetic make-up of ni-Vanuatu, we collected >4,000 blood samples from contemporary individuals between 2003 and 2005, and genotyped a selection of 1,433 of these sampled individuals at 2,358,955 single nucleotide polymorphisms (SNPs). After merging with 179 high-coverage whole genome sequencing data [18] and excluding 173 low-quality, duplicated or related samples, a total of 1,439 ni-Vanuatu samples was included in subsequent analyses, including those from 522 males and 917 females living in 29 islands and 179 different villages (Fig. 1A and Table S1).

**Figure 1.**
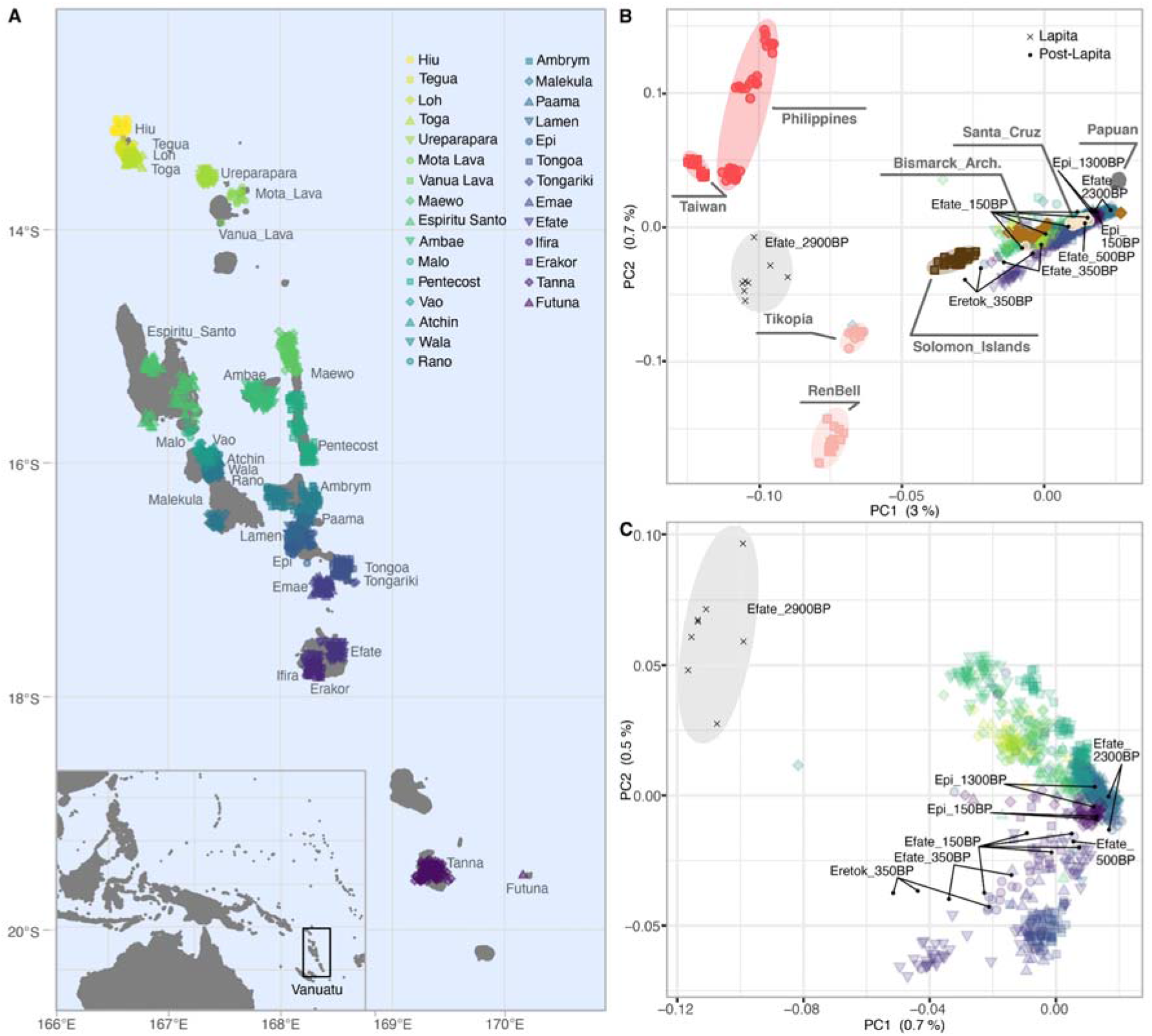
Sampling locations and genetic structure in Vanuatu. (A) Map showing the sampling location of the 1,439 contemporary ni-Vanuatu individuals. The inset at the bottom left shows the location of Vanuatu in the Asia-Pacific region. Noise was added to sampling locations to facilitate visualization. (B,C) Principal Component Analysis (PCA) of genotypes of ni-Vanuatu at 301,774 SNPs, in the context of (B) the broad Asia-Pacific region and (C) the Vanuatu archipelago only. (A-C) Each point indicates an individual, colored according to the latitude of their island of residence. (B,C) Black crosses and points indicate projected ancient samples from Vanuatu dated to the Lapita and Post-Lapita periods, respectively.

Principal Component (PCA) and ADMIXTURE analyses indicate that contemporary ni-Vanuatu fall on a genetic gradient between East Asian-related and Papuan-related populations (Fig. 1B and Fig. S1), supporting the view that their ancestry derives from these two population groups. When projecting ancient samples from Vanuatu, we found that Lapita individuals show higher affinity with present-day East Asian-related populations, whereas Post-Lapita individuals are closer to Papuan-related populations, in line with a Papuan-related population turnover occurring after the Lapita period [8–10, 13]. Furthermore, contemporary ni-Vanuatu show high genetic similarities with individuals from the Bismarck Archipelago, in line with the hypothesis that their Papuan-related ancestors originated from these islands [8–10, 13, 18]. However, the extensive geographic coverage of the dataset presented here enabled us to reveal substantial genetic substructure among ni-Vanuatu islanders. PCA showed that genetic variation in ancient and contemporary individuals from Vanuatu is explained by two contiguous but distinct groups that broadly reflect geography (Fig. 1C and Fig. S2). When considering the number of ancestral components that is most supported by ADMIXTURE (*K*_ADM_ = 9), populations from different islands also show different ancestral components (Fig. S1). These results reveal that contemporary ni-Vanuatu show substantial genetic differentiation (Fig. S3), which could result either from different admixture histories during the settlement period and/or from the existence of barriers to gene flow that formed after the settlement of the archipelago.

To gain further insights into the genetic history of ni-Vanuatu, we next assessed the fine-scale genetic structure of the 1,439 sampled individuals, using ChromoPainter and fineSTRUCTURE [19–21]. Haplotype-based clustering revealed a first separation between ni-Vanuatu living north and south of the strait separating Epi and Tongoa islands (*K*_FS_ = 2; Fig. S4). At *K*_FS_ = 4, populations separate into clusters, here referred to as the *Banks and Torres Islands* cluster and the *North East, North West* and *Central-South Vanuatu* clusters (Fig. 2A and Fig. S4 and Table S2), that are in general agreement with the classification of Oceanic languages spoken in the archipelago [22–24]. Of note, individuals from the north and south of Pentecost island cluster with individuals from different neighboring islands, indicating that the sea does not necessarily act as a barrier to gene flow in the region, in line with linguistic and ethnographic data that reveal cultural networks between islands [22, 23, 25]. Likewise, in the Ambae island, northwest and south inhabitants cluster separately (Fig. 2A), suggesting that the periodic volcanic activity of the Ambae caldera has affected gene flow in the region. At *K*_FS_ = 20, i.e., the highest *K*_FS_ value for which statistical robustness remains maximal (Methods) [20], we found that genetic clusters are often island-specific (Fig. 2A, Figs. S4-S7 and Table S1). These observations suggest that clusters inferred by fineSTRUCTURE are reliable, as they reflect expected geographic and linguistic barriers to gene flow.

**Figure 2.**
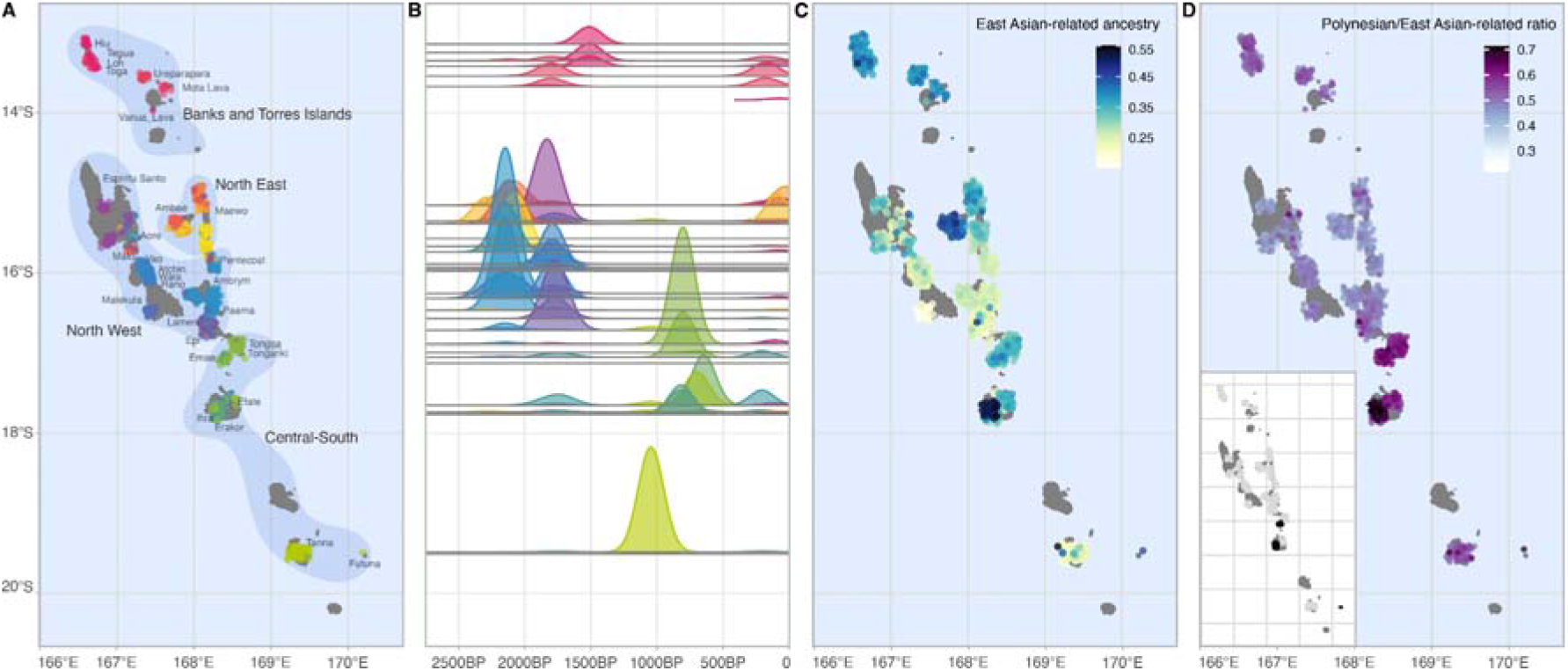
Fine-scale genetic structure and admixture in Vanuatu. (A) Map showing the clustering of 979 ni-Vanuatu into 20 genetic clusters, according to fineSTRUCTURE (*K*_FS_ = 20). Each point indicates an individual, located according to their village of residence. Colors indicate genetic clusters, so that the closer the colors, the closer the clusters. Noise was added to sampling locations to facilitate visualization. (B) Admixture date estimates for each genetic cluster, based on 100 bootstrap replicates. Colors are the same as those used in (A). The x-axis shows the admixture date in years before present, assuming a generation time of 28 years. The y-axis indicates the latitude of the islands assigned to each genetic cluster. The heights of the density curves are proportional to the sample size of each cluster. (C) Proportion of East Asian-related ancestry estimated by SOURCEFIND. (D) Ratio of the Polynesian and East Asian-related ancestry proportions, estimated by SOURCEFIND. The inset at the bottom left shows the location of Polynesian-speaking individuals, indicated by black points. (C-D) Each point indicates an individual, colored according to their ancestry proportions.

### The Post-Lapita Shift to Papuan-related Ancestry was Geographically Uneven

We detected important differences among fineSTRUCTURE clusters in the proportions of East Asian-related ancestry, that is, ancestry related to contemporary populations from either Taiwan and the Philippines or Polynesia (range of SOURCEFIND estimates: 0.15 to 0.55; Fig. 2C, Figs. S8-S11 and Table S3). East Asian-related ancestry was found to be lowest for the *North Vanuatu* clusters (e.g., Malekula, Ambrym, Epi; median = 0.230, SD = 0.056) and highest in the *Central-South Vanuatu* cluster (i.e., Mele and Imere in Efate island, Makatea, Tongamea and Vaitini in Emae island, and Tongoa and Ifira islands; median = 0.323, SD = 0.077), where “Polynesian outlier” communities live today [11, 12]. Nevertheless, East Asian-related ancestry is also high in islands where Polynesian ancestry is low, such as Ambae (median = 0.405, SD = 0.042; Fig. 2D), suggesting that differences in East Asian-related ancestry are not due solely to differences in Polynesian ancestry (Figs. S10 and S11 and Table S3).

To assess whether differences in East Asian-related ancestry originate from a geographically uneven ancestry turnover or separate admixture events between distinct populations, we dated admixture in each genetic cluster separately. Besides more recent events relating to Polynesian migrations (see next section), all estimates overlap the same time period that ranges from 1,700 to 2,300 ya (Fig. 2B, Fig. S12 and Table S4), suggesting that all ni-Vanuatu share the same admixture history. To test this hypothesis more formally, we evaluated if the Papuan-related ancestry carried by present-day ni-Vanuatu derives from a single source. PCA and *f*_4_-statistics of the form *f*_4_(*X*, New Guinean highlanders; Solomon or Bismarck Archipelago islanders, East Asians) indicate that all ni-Vanuatu show similar genetic relatedness with populations from the Bismarck Archipelago, New Guinea or the Solomon islands (Figs. S13 and S14), in agreement with aDNA data [13]. Furthermore, the haplotype-based SOURCEFIND method detected that the same cluster of Bismarck Archipelago islanders is the source that contributed the most to all ni-Vanuatu (Fig. S15 and Table S5). Of note, SOURCEFIND also detected a small contribution from New Guinean highlanders and Santa Cruz islanders in all ni-Vanuatu clusters, which negatively correlates with Polynesian ancestry (*r* = −0.378, *P*-value < 2.22×10^-16^), probably because Polynesian source populations capture some Papuan-related ancestry. Collectively, these findings indicate that ni-Vanuatu are descended from relatively synchronous admixture events between the same sources of Papuan- and East Asian-related ancestry; yet, their proportions differ markedly across islands, suggesting that the dramatic Papuan-related ancestry shift that started ~2,500 ya was geographically uneven.

### Polynesian Migrations Did Not Necessarily Trigger Language Shifts

Admixture proportions and date estimates indicate that ni-Vanuatu ancestry partly derives from a third and more recent migration originating from Polynesia (Fig. 2B-D and Tables S3 and S4). We dated admixture events between 600 and 1,000 ya for genetic clusters predominant in Mele and Imere (Efate island), Makatea, Tongamea and Vaitini (Emae island), as well as in Ifiraand Futuna islands (Fig. 2B, Fig. S12 and Table S4), where Polynesian languages are spoken today [12, 26]. These clusters show higher Polynesian ancestry and, therefore, higher East Asian-related ancestry proportions, relative to *Banks and Torres Islands* and *North Vanuatu* clusters (SOURCEFIND estimates; Fig. 2C-D, Figs. S8-S11 and Table S3). Ni-Vanuatu Polynesian speakers show a higher ratio of Polynesian-to-East Asian ancestry, when compared to non-Polynesian speakers (ratio = 0.647 *vs*. 0.479; Wilcoxon test *P*-value < 2.22×10^-16^; Fig. 2D and Fig. S10). Furthermore, *f*_4_-statistics of the form *f*_4_(*X*, East Asians; Tongans, New Guinean highlanders) suggest that ni-Vanuatu from Efate, Ifira and Emae share more alleles with Polynesians than other ni-Vanuatu do (Fig. S16).

Interestingly, our analyses also revealed that Polynesian ancestry is not restricted to Vanuatu islands where Polynesian languages are spoken. Non-Polynesian-speaking populations assigned to the *Central-South Vanuatu* (e.g., Tongoa, Tongariki and Tanna) and the *Banks and Torres islands* clusters also show a higher Polynesian-to-East Asian ancestry ratio, relative to *North East* and *North West* clusters (ratio = 0.557 and 0.510, *vs*. 0.461; Wilcoxon test *P*-value < 2.22×10^-16^; Fig. 2D, Figs. S10 and S11 and Table S3). Furthermore, estimated admixture dates are similar among groups from the *Central-South Vanuatu* cluster that speak or not Polynesian languages (Fig. 2B, Fig. S12 and Table S4). These results support the view that Polynesian migrations, as well as subsequent contacts between Vanuatu islands [12, 27] or with “Polynesian outliers” from the Solomon islands [28], also introduced Polynesian ancestry among non-Polynesian-speaking groups.

Conversely, we found no evidence of admixture with Polynesians in individuals assigned to *North Vanuatu* clusters, including Epi islanders, despite the geographic proximity between Epi and Tongoa (Fig. 2D, Figs. S10 and S11 and Table S3). Of note, ADMIXTURE, PCA and fineSTRUCTURE analyses separate ni-Vanuatu into northern and southern populations, the frontier between the two being located between Epi and Tongoa (Figs. 1C and 2A and Fig. S1). The strait that separates the two islands today is the location of the Kuwae caldera [29, 30], whose volcanic activity may have been a barrier to gene flow and/or may have triggered large-scale population movements that disrupted the isolation by distance patterns. Together, these results reveal that, since 1,000 ya onward, Polynesians migrated to Vanuatu where they admixed with local populations, and that such interactions did not necessarily result in a shift to Polynesian languages.

### A Limited Genetic Impact of European Colonization

Our genetic data indicate that admixture between ni-Vanuatu and Europeans has been rare or has left few descendants in Vanuatu. In total, only 28 individuals out of 1,439 (1.95%) show a genetic contribution from Europeans higher than 1%, according to SOURCEFIND analyses (range: 0.03% to 35.3%; Fig. S8 and Table S3). The two fineSTRUCTURE clusters with the highest European ancestry (median = 0.093 and 0.107, *vs*. 0.002 for other clusters) show an admixture event in the last 120 years (mean = 78 ya, 95%CI: [74–81] and mean = 111 ya, 95%CI: [104-118]) (Fig. 2B and Table S4). Similarly, the three other clusters that include individuals carrying more than 1% of European ancestry show a pulse of admixture occurring in the last 200 years. These results are consistent with historical records; early, fleeting European contacts with ni-Vanuatu began in 1606, with subsequent documented exploratory voyages in 1768, 1774 and 1809; yet, it was only from around 1829 onwards that contacts became more common, when Christian missionaries and European colonists settled in the archipelago and the first intermarriages were reported [31, 32].

### Genetic Admixture in Vanuatu was Sex-Biased

Genetic studies have suggested that Papuan-related migrations from the Bismarck Archipelago into Remote Oceania were male-biased, because contemporary Polynesians and ancient individuals from present-day Vanuatu show lower Papuan-related ancestry on the X chromosome, relative to autosomes [8, 13]. To confirm that admixture between the ancestors of present-day ni-Vanuatu was sex-biased, we estimated Papuan- and East Asian-related ancestry in ni-Vanuatu on each chromosome separately, using local ancestry inference [33]. We found that Papuan-related ancestry is indeed significantly lower on the X chromosome, relative to autosomes (*α_X_* = 75.2% *vs*. *α*_auto_ = 79.8%, Wilcoxon test *P*-value < 1.36×10^-5^; Fig. 3A), in agreement with aDNA results [13]. These values were similar between Polynesian and non-Polynesian speakers (Fig. S17A, Wilcoxon test *P*-value = 0.13), which indicates that it is not explained by recent migrations from Polynesia. We replicated the results using high-coverage genome sequences from a subset of 179 ni-Vanuatu [18], implying that our results are not biased due to SNP ascertainment (Fig. S17B). Assuming that admixture proportions have reached equilibrium values [34], we estimated that the genetic contribution of Papuan-related males to ni-Vanuatu was 27% higher than that of Papuan-related females (*α*_m_ = 93.5% *vs*. *α*_f_ = 66.1%). Accordingly, Y chromosomes of ni-Vanuatu are dominated by haplogroups found at high frequency in Near Oceanians (e.g., M1a3b2, S1), whereas mitochondrial DNAs (mtDNA) show a high proportion of haplogroups typically found in East Asia (e.g., B4a1a1, E1a2a4) (Fig. 3B and Table S1). Collectively, these results support the notion that ni-Vanuatu ancestry predominantly results from admixture between Papuan-related males and East Asian-related females.

**Figure 3.**
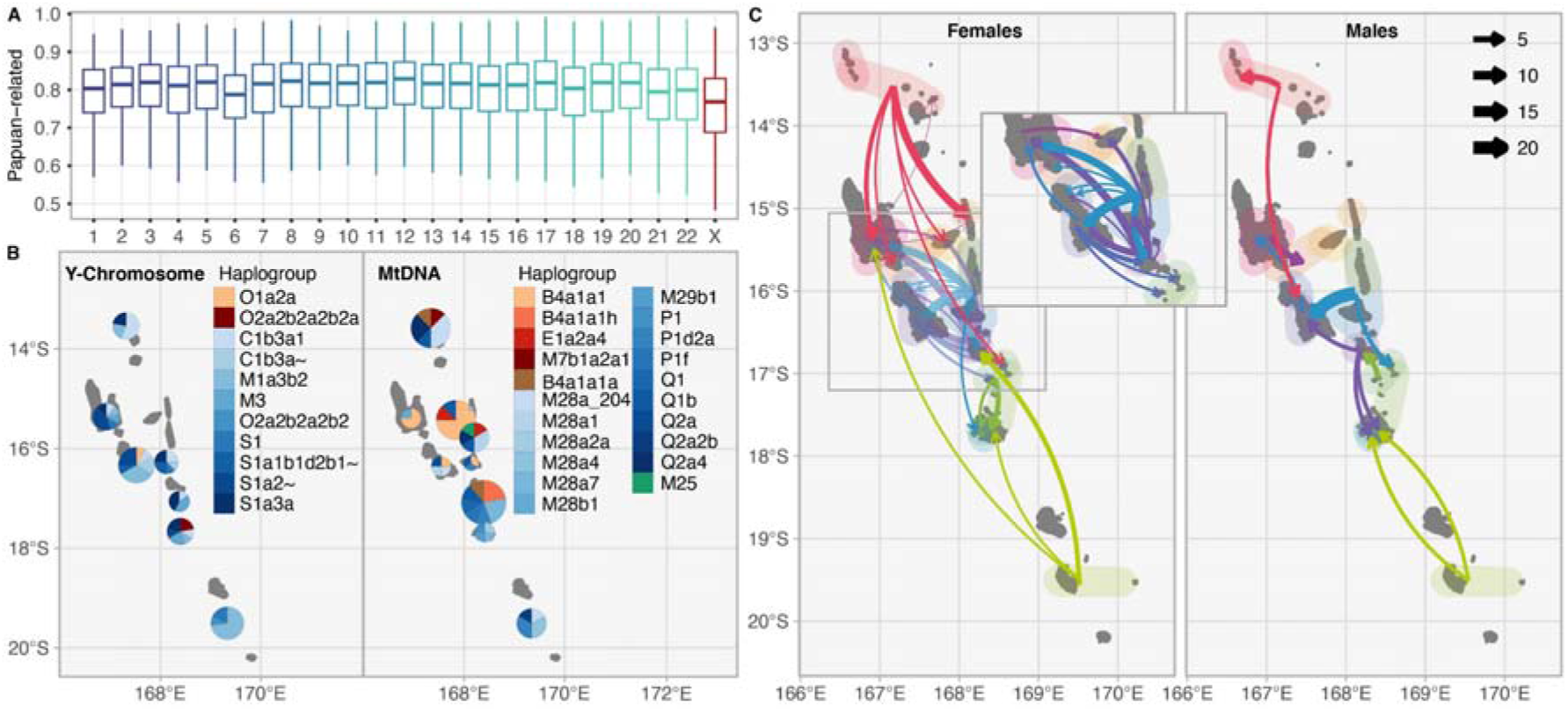
Sex-biased admixture and migration patterns in Vanuatu. (A) Papuan-related ancestry proportions in ni-Vanuatu, estimated for the 22 autosomes and the X chromosome separately by RFMix. (B) Frequencies of Y chromosome and mtDNA haplogroups, colored according to their assumed origins (i.e., shades of blue or red indicate Papuan- or East Asian-related origins, respectively). Haplogroups were inferred from high-coverage genome sequencing data obtained for a subset of 179 ni-Vanuatu [18]. (C) Recent migrations among Vanuatu islands, inferred based on fineSTRUCTURE clusters. The arrows connect the location of the genetic cluster to which individuals were assigned to their actual place of residence. The colors indicate the predominant genetic cluster in the island of origin. The width is proportional to the number of inferred migrant individuals, relative to the number of females or males in the genetic cluster.

### Recent Migrations are Influenced by Residence Rules and Urbanization

The genetic structure of contemporary ni-Vanuatu is also expected to reflect socio-cultural practices (e.g., social networks, exchange and marriage rules) that have culturally evolved since the settlement of the archipelago. We leveraged the high-resolution genetic data to infer recent migrations between Vanuatu islands – indicated by individuals who inhabit an island but belong to a genetic cluster that is prevalent in another island – and determine if these migrations have involved mainly females or males, in line with virilocal or uxorilocal postmarital residence rules, respectively. We found that 5.70% of the sampled individuals (54 individuals) migrated at a large geographical scale (*K*_FS_ = 4), while 11.81% (112 individuals) migrated at a local scale (*K*_FS_ = 20; Table S6), suggesting more genetic connections between closer islands. We estimated that local mobility among females is higher than among males (*K*_FS_ = 20; odds ratio = 2.20, Fisher’s exact test *P*-value = 9.16×10^-4^; Fig. 3C), as expected under virilocal residence and/or female exogamy [35]. The same trend was observed at a larger geographical scale, when considering migrations between four broader regions (*K*_FS_ = 4; odds ratio = 1.86, *P*-value = 0.075) and when restricting the analyses to the reported birth place of male-female couples included in the dataset (Fig. S18) (odds ratio = 1.96, *P*-value = 6.89×10^-3^). Notably, comparisons of the places of birth and residence of sampled individuals did not support female-biased migrations (odds ratio = 0.807, *P*-value = 0.18; Fig. S19 and Table S6), possibly reflecting a bias in the self-reported birth places. This might occur if women are inclined to report their husband’s or children’s place rather than their father or mother’s place [36] or under other, more complex patterns.

We also explored the direction of the inferred migrations, and found that both female and male migrations mainly occurred from the northernmost and southernmost islands towards the center of the archipelago (Fig. 3C), where Port Vila - the largest city - has developed [37]. Migrations are also common among North Vanuatu islands, consistent with a long-term network of cultural and material exchanges in the region [25, 28]. Thus, our genomic data reflect the migration patterns that characterize the recent history of ni-Vanuatu, including residence rules and urbanization.

### No Evidence for Endogamous Practices among Ni-Vanuatu Spouses

Human kinship systems vary tremendously, regulating marriage, exchange, endogamy or exogamy according to relational concepts [38]. Given the small census size of ni-Vanuatu populations, it is interesting to consider whether and how sociocultural features like marriage and exchange rules influence local levels of genetic relatedness. We found that genetic relatedness was higher within islands than between islands (Wilcoxon test *P*-value < 2.22×10^-16^; Figs. S20-S22 and Table S7), consistent with an isolation by distance model (Mantel test *P*-value = 0.001; 1,000 permutations). Accordingly, out of the 287 male-female couples included in the dataset, 78.4% were born on the same island (Fig. S23). Genetic relatedness is also higher among inhabitants of the same village, relative to individuals living on the same island (Wilcoxon test *P*-value < 2.22×10^-16^) (Fig. S20 and Table S7), suggesting that the local community is often the source of marriage partners. However, despite the generally high levels of genetic relatedness observed between ni-Vanuatu, we did not observe an excess of genetic relatedness among couples (Figs. 4A, S23 and S24). Specifically, spouses tend to show slightly lower kinship coefficients than random pairs of individuals from the same village, when excluding first-degree related individuals from all possible pairs (logistic regression model *P*-value = 0.057; resampling *P*-value = 0.079) (Figs. S24 and S25) and when adjusting on differences in ancestry (*P*-value < 0.05; Figs. 4A). Assuming that our total sample is not biased towards related individuals, these results suggest that endogamy is not a common practice among contemporary ni-Vanuatu and more generally illustrate that populations can show high levels of genetic relatedness in the absence of endogamous marriage practices.

**Figure 4.**
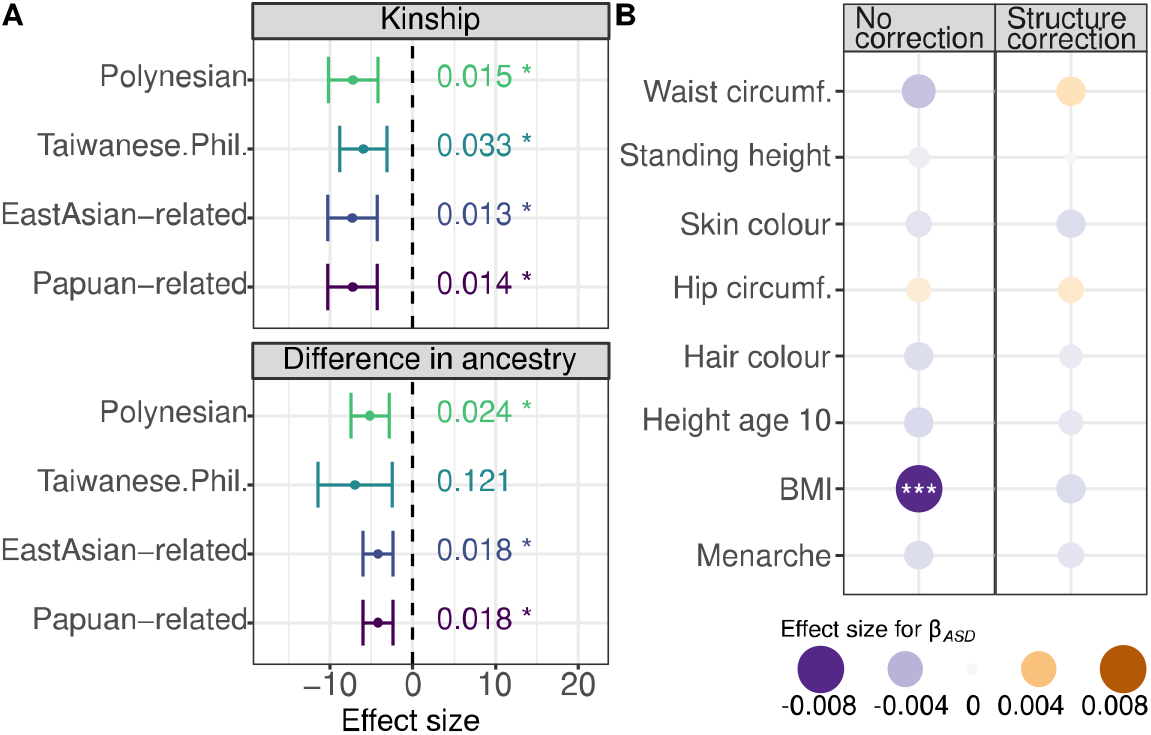
Ancestry- and trait-based assortative mating in Vanuatu. (A) Effects of kinship and ancestry differences on partner choice. Effect sizes were estimated with a logistic regression model, while accounting for population structure (island of birth and village of residence). The effect size and *P*-value were estimated using different ancestries as predictors, independently. (B) Increased (purple) or decreased (orange) genetic similarity among spouses (as measured by 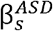) at trait-associated SNPs, relative to non-associated SNPs. Results on the left are based on a logistic model that includes only the genotype dissimilarity among spouses at each SNP, whereas results on the right are based on the logistic model that also controls for possible confounders (i.e. geography, kinship and ancestry differences).

### Ni-Vanuatu Spouses Tend to Share Similar Genetic Ancestry

Research on admixed populations from other parts of the world has shown that, in addition to low genetic relatedness, partners tend to show similar genetic ancestry [16], because mating often occurs within socio-cultural groups, which can correlate, in turn, with genetic ancestry. To verify whether this phenomenon is observed in Vanuatu, we implemented a logistic regression model that jointly estimates the effects of geography, genetic relatedness and genetic ancestry on the probability to be partners (Methods). These analyses confirmed that spouses tend to originate from the same island, while showing lower kinship coefficients than non-spouses (Fig. 4A). Importantly, we found that spouses show lower differences in genetic ancestry than non-spouses, when considering Papuan-related (β = −4.233, *P*-value = 0.018), East Asian-related (β = −4.221 *P*-value = 0.018) or Polynesian (β = −5.197, *P*-value = 0.024) ancestry. These results suggest that ni-Vanuatu tend to mate with a partner who carries similar genetic ancestry.

Two hypotheses have been proposed to explain these observations. First, social assortative mating may underlie the observed signal, as genetic ancestry can correlate with socio-cultural structure. Second, spouses may choose their partner because they share biological traits, such as physical appearance [39, 40]. To test these hypotheses, we searched for genomic loci that could play a role in partner choice, by including, in the logistic regression model, a term that measures dissimilarity between individuals at each SNP (Methods). No SNP showed statistical evidence for a significantly lower or higher genotype similarity between spouses, when accounting for multiple testing (Fig. S26). To test for assortative mating according to polygenic traits, we then evaluated if genotype similarity between spouses is significantly higher or lower at SNPs associated with candidate traits relating to physical appearance, when compared to non-associated SNPs. When we did not account for genetic structure and ancestry-associated assortative mating (i.e., effect sizes are estimated from a model where these confounders are not included, see Methods), we found evidence for assortative mating according to body mass index (BMI, Fig. 4B). However, when accounting for such possible confounders, we found no statistical evidence that genotypes at trait-associated variants are more similar or dissimilar between spouses than expected. Collectively, our results do not support a marked tendency for partner choice according to genetic or phenotypic features, and suggest instead the occurrence of assortative mating driven by social structure as the cause for ancestry-based assortments among ni-Vanuatu.

## DISCUSSION

By leveraging an extensive genomic dataset of 1,439 contemporary individuals, we show here that ni-Vanuatu initially descend from admixture between the same ancestral populations: an East Asian-related population, which shares genetic affinities with groups living today in Taiwan and the Philippines, and a Papuan-related population, which shares genetic affinities with groups living today in the Bismarck Archipelago. We also estimate that admixture occurred after the Lapita period, ~1,700-2,300 years ago, and was relatively synchronous across islands, in agreement with a peopling history common to all the archipelago. Thus, our results suggest that the high cultural diversity of ni-Vanuatu results from a rapid cultural diversification that developed *in situ*, as suggested by linguistic, archaeological and archaeogenetic studies [13, 14, 28]. Nevertheless, our analyses cannot definitely rule out that, after the Lapita period, Vanuatu islands were settled by different, already admixed groups carrying varying levels of East Asian-related and Papuan-related ancestry, as recently suggested [18]. Furthermore, we caution that admixture date estimates from modern DNA data are uncertain and can be biased downward when admixture was gradual [41], which was likely the case in Vanuatu [10]. Additional aDNA time transects from multiple islands will be required to provide a definitive picture of the admixture history of ni-Vanuatu.

Our analyses reveal substantial differences in East Asian-related ancestry proportions between islands. We show that these differences do not result solely from Polynesian migrations, as our haplotype-based analyses could differentiate ancestry attributed to the Austronesian expansion from that introduced by Polynesians. A compelling example is Ambae, where East Asian-related ancestry is 1.8-times higher than in surrounding islands but where Polynesian ancestry is low. Assuming a simple admixture model, these findings imply that the major population turnover following the arrival of Papuan-related peoples was geographically uneven, possibly because, at the time of admixture, the two ancestral populations of ni-Vanuatu were of different sizes across islands, some of which being preferentially settled by East Asian-related groups and others by Papuan-related groups. Lastly, we confirm that admixture was sex-biased [8, 13]; either Papuan-related migrants were predominantly males or both males and females migrated, but admixture was more common between Papuan-related males and East Asian-related females.

A recent archaeogenetic study has reported that ancient ni-Vanuatu from the Chief Roi Mata’s Domain in Efate show genetic similarities with Polynesians [13], supporting the occurrence of migrations from Polynesia, which were previously postulated by linguistic studies [11, 12]. We confirm that “Polynesian outlier” communities in Vanuatu are descended from admixture events between Polynesians and local populations. We dated these admixture events to 600-1,000 ya, in line with archaeological records [11]. Furthermore, we extend previous findings by mapping the genetic impact of Polynesian migrations to some Vanuatu islands where Polynesian languages are not spoken today (e.g., Makura, Tongoa, Tongariki and Tanna) [12]. These results indicate that genetic interactions between ni-Vanuatu and Polynesian incomers did not systematically trigger shifts to Polynesian languages. Intriguingly, Polynesian ancestry is not detected north of the Kuwae caldera, a large submarine volcano that separates Tongoa and Epi islands. Geological data have shown that the Kuwae volcano erupted in ca. 1452, producing among the largest volumes of magma and aerosol ever recorded [29, 30], and oral traditions and linguistic evidence suggest that, after this eruption, Tongoa and Epi were repopulated by distant populations [42, 43]. Our genetic results support this view; present-day Tongoa and Emae islanders, as well as Epi and southwest Malekula islanders, show close genetic affinities (Fig. S4), at odds with the gradual genetic differences expected under isolation by distance. Nowadays, ni-Vanuatu living north and south of Kuwae form two genetic groups with distinct socio-cultural practices (e.g., grade-taking and chiefly title political systems) [44], indicating that the area has remained a genetic and cultural frontier.

By building upon the well-defined genetic history of Vanuatu, we also explored how genetic diversity has been shaped by cultural practices. While levels of genetic relatedness are high among ni-Vanuatu, we do not find evidence for generalized endogamy, which challenges the frequent association geneticists make between the two processes [17, 45–47]. Nonetheless, even if ni-Vanuatu spouses are generally less related than non-spouses, we show that their genetic ancestries are more similar than expected, indicating that mating in Vanuatu is not random. Importantly, ancestry similarity between partners is not stronger at trait-associated SNPs, suggesting that ancestry-associated assortments are due to social structure, which may, in turn, be correlated with levels of East Asian-related and/or Polynesian ancestry. Other studies have suggested non-random mating according to ancestry in regions of the world where socio-cultural structure is highly correlated with ancestry [16, 48]. Our findings extend the occurrence of such socio-cultural assortments to Oceanians, raising questions of how common this phenomenon is in human societies and whether non-random mating should systematically be accounted for in human genetic studies. Collectively, our study emphasizes the need to include diverse populations in genetic studies, not only to address key anthropological and evolutionary questions that are important for specific geographic regions, but also to identify factors shaping the genetic diversity of human populations as a whole.

## METHODS

### RESOURCE AVAILABILITY

#### Lead Contact

Further information and requests for resources and reagents should be directed to and will be fulfilled by the Lead Contact, Etienne Patin.

#### Materials Availability

This study did not generate new unique reagents.

#### Data and Code Availability

The SNP array data generated in this study has been deposited to the European Genome-Phenome Archive (EGA) under accession number EGAS00001005910.

### EXPERIMENTAL MODEL AND PARTICIPANT DETAILS

The sampling survey was conducted in the Republic of Vanuatu between April 2003 and August 2005. The purpose of the study was the estimation of the seroprevalence of HTLV-1 viral infection and the assessment of human genetic diversity in ni-Vanuatu. The recruitment of participants was carried out after the agreement of the Ministry of Health of Vanuatu, the head of each Directorate from the sampled province, and the chief of the sampled village. The data collectors, Olivier Cassar, Helene Walter, Woreka Mera and Antoine Gessain, were accompanied by the village chief and/or the head of the local dispensary. The nature and the scope of the study were explained in detail by Olivier Cassar in English, and by Helene Walter in Bislama (i.e., an English-based pidgin-creole [14] that is the main *lingua franca* of Vanuatu), during information meetings organized in each village. Participants could ask any questions after the information meeting. After several hours of reflection, each volunteer participant of at least 18 years of age was asked to sign a written informed consent form, including consent for research on human genetic diversity. Sex, age and birth place, as well as the date and place of blood collection, were collected through a structured questionnaire. Couples were identified through interviews in English or Bislama, and were preferentially sampled. Blood samples were collected either at the local dispensary, a gymnasium or a hut provided by the village chiefs.

The study received approval from the Institutional Review Board of Institut Pasteur (n°2016-02/IRB/5) and the Ministry of Health of the Republic of Vanuatu. It was conducted in full respect of the legal and ethical requirements and guidelines for good clinical practice, in accordance with national and international rules. Namely, research was conducted in accordance with: (i) ethical principles set forth in the Declaration of Helsinki (Version: Fortaleza October 2013), (ii) European directives 2001/20/CE and 2005/28/CE, (iii) principles promulgated in the UNESCO International Declaration on Human Genetic Data, (iv) principles promulgated in the Universal Declaration on the Human Genome and Human Rights, (v) the principle of respect for human dignity and the principles of non-exploitation, non-discrimination and non-instrumentalisation, (vi) the principle of individual autonomy, (vii) the principle of justice, namely with regard to the improvement and protection of health and (viii) the principle of proportionality. The rights and welfare of the participants have been respected, and the hazards did not outweigh the benefits of the study. Feedback to local communities and partners in Vanuatu is planned for 2023, under the guidance of H.C. The results of this study will be presented to key stakeholders, including the Ministry of Health and the Vanuatu Cultural Center (Port Vila). Written and recorded resources will be provided in Bislama and distributed to interested individuals and communities.

### METHOD DETAILS

#### SNP genotyping and quality filters

Five millilitres of blood were obtained by venepuncture from each volunteer participant and transferred to the Institut Pasteur of New Caledonia, where plasma and buffy coats were isolated, frozen, and stored at −80°C. Samples were then transferred to the Institut Pasteur in Paris (France). Out of the 4,428 collected samples, 1,433 samples were selected for genetic analyses. These samples were selected in order to (i) cover the largest number of islands and villages, (ii) cover locations where “Polynesian outliers” exist nowadays and (iii) include as many couples as possible. After sample selection, a total of 179 different villages were covered, located on 29 islands (Table S1). DNA was purified from frozen buffy coats at the Institut Pasteur of Paris (France), using QIAamp DNA Blood Mini Kit protocol, and eluted in AE buffer. DNA concentration was quantified with the Invitrogen Qubit 3 Fluorometer using the Qubit dsDNA broad-range assay. Prior to SNP array genotyping, DNA integrity was checked on agarose gels.

The 1,433 selected samples were genotyped on the Illumina Infinium Omni 2.5-8 array (San Diego, California). Genotype calling was performed using the Illumina GenomeStudio software. We excluded 2,491 SNPs with missing annotations, 9,661 duplicated SNPs, 1,772 SNPs with a GenTrain score < 0.4, SNPs with a missingness > 0.05 and SNPs that deviate from Hardy-Weinberg equilibrium (i.e., *P*-value < 0.01 in more than one Vanuatu island). Only autosomal SNPs were kept for the analyses, unless otherwise stated. After filters, a total of 2,269,868 SNPs were kept. When the analyses required minor allele frequency (MAF) filters, a MAF > 0.01 threshold was applied. When a linkage disequilibrium pruning was required, we pruned the data using a window size of 50 Kb, a step size of 5 SNPs and a *r^2^* threshold of 0.5. After all these filtering steps were applied, the remaining number of SNPs was 294,806. All filters were applied using PLINK v.1.9b [49].

#### Sample quality filters

The highest genotype missingness per sample was 0.018. We removed 21 samples with outlier values for heterozygosity (mean ± 3 SD), suggestive of DNA contamination, leaving 1,412 samples. Cryptic relatedness between samples was detected using KING v.2.1 [50]. We excluded 456 samples with a kinship coefficient > 0.08 with another sample, whenever analyses required unrelated samples. For 12 samples, the reported sex did not match the genetic sex inferred by the Y- and X-chromosome call rates. These include one couple, for which the reported sex is the opposite to the genetic sex. We thus exchanged the sex of the two samples, as it is most likely an error on the sample annotation. We did not include the 10 remaining samples, when performing analyses relating to mating practices and sex-specific migrations. After quality filters, a total of 287 couples was present in the dataset.

#### Demographic information

Demographic data include the island of birth and the village of residence. Birth place was missing for 73 samples and the residence for 49 samples. From this information, we retrieved, for each sample, the geographical coordinates of the birth place (around the center of the island) and of the place of residence (around the village of residence), using MapAction reference maps (www.mapaction.org). We considered as the locations of “Polynesian outliers” all the villages within Futuna, the villages of Mele and Imere in Efate, Ifira island and the villages of Makatea, Tongamea and Vaitini in Emae [12]. Among the 287 couples, 207 reported the same island of birth, while 57 reported a different island. This information was missing for 23 couples.

#### Merging with reference datasets

We merged the new SNP array data with whole genome sequences of different populations across the Pacific [18] and worldwide populations from the SGDP project [51]. We also merged the dataset with SNP array data from Oceanian populations [41], to perform some population genetics analyses. Datasets were merged using PLINK v.1.9b [49]. Transversions were excluded, to avoid allele strand inconsistences.

#### Haplotype phasing

We phased the dataset using SHAPEIT v.2 [52, 53] using 500 conditioning states, 10 burn-in steps, 50 Markov chain Monte Carlo (MCMC) main steps, a window length of 1 cM and an effective population size of 15,000. We used 1000 Genomes data [54] as a reference panel and therefore removed all array SNPs that did not align with this data, prior to haplotype phasing.

#### Genetic structure: PCA and ADMIXTURE

We performed Principal Component Analyses (PCA) with the ‘SmartPCA’ algorithm implemented in EIGENSOFT v. 6.1.4 [55]. aDNA samples were projected on PCs by using the “lsqproject” and “shrinkmode” options. We inferred population structure with ADMIXTURE v. 1.3.0 [56], using 10 different seeds and assuming that *K*_ADM_ varies from 2 to 17, and visualized ADMIXTURE results using pong v1.4.7 [57]. We estimated pair-wise *F*_ST_ values using the StAMPP R package [58], and assessed their significance with 100 bootstraps. We computed *f*_4_-statistics with ADMIXTOOLS v.6.0 and estimated standard errors by block jackknife.

#### Genetic structure: ChromoPainter and fineSTRUCTURE

We ran ChromoPainter [19] to infer haplotype sharing among individuals. This algorithm is based on a Hidden Markov Model in which each sample is treated as a “recipient”, i.e., a mosaic of haplotypes from a set of “donor” samples. We performed different analyses with ChromoPainter. In Analysis 1, we inferred the genetic structure of ni-Vanuatu running ChromoPainter only for the unrelated ni-Vanuatu samples. We set each sample both as a recipient and as a donor for all the samples (-a mode). We first estimated the global mutation probability and the switch rate with 10 expectation-maximization (EM) iterations, by using the ‘-in -iM’ options and by averaging estimates obtained for chromosomes 1, 7, 13 and 19. The estimated values were 1.542×10^-4^ and 81.415, respectively. Then, we ran ChromoPainter in the same mode for all chromosomes but fixing the parameters to estimated values. In Analysis 2, we repeated the same process independently for the reference populations. In this case, we ran ChromoPainter only for the samples outside Vanuatu [18, 51] and repeated the two-step process described above. In this case, the global mutation probability was estimated to 4.53×10^-4^ and the switch rate was 255.42.

We then produced a coancestry matrix by summing the coancestry matrices for all chromosomes and estimated the *C* parameter with ChromoCombine. The *C* factor was estimated at 0.452 for the Vanuatu dataset (Analysis 1), and 0.498 for the reference population dataset (Analysis 2). We applied the model-based Bayesian clustering method fineSTRUCTURE v.2.0.7 [19] on ChromoPainter coancestry matrices. We ran 1 million burn-in iterations (-y option) and 1 million MCMC iterations (-x option) and sampled values every 10,000 iterations. We then inferred the fineSTRUCTURE tree (-m T option). We repeated this process 5 times, with different random seeds.

Following a previous study [20], we estimated for each individual the robustness of the clustering assignation, by comparing the final state (i.e., the state with the highest posterior probability) with the 100 MCMC samples. For each individual *i*, we estimated 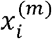, the number of individuals clustering with *i* for each of the *m* = 1,…,100 MCMC samples. We also estimated 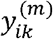, the number of individuals that cluster with *i* in the final state and in each of the MCMC samples, for each inferred cluster *k*. Finally, we computed 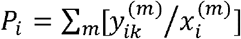. Therefore, we ended up with a matrix of this sum *P_i_*, for all individuals and for each genetic cluster, that shows the robustness in the assignation of each individual to its corresponding cluster in the final state. We repeated this process for *k* = 2,…,*K*, *K* being the maximum number of clusters identified by fineSTRUCTURE. We also repeated the process for the 5 random seeds and compared the performance of each seed. Fig. S6 shows the number of ni-Vanuatu individuals with *P_i_* < 0.9 for each seed *s* and for each *k* = 2,…*K*. These estimations allowed to test the robustness in the cluster assignation of each individual, to compare the performance across seeds and to determine the robustness at each *k* value.

To compare robustness among seeds, we followed a similar approach. We estimated 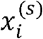, where *s* = 1,…,5, which corresponds to the number of individuals clustering with individual *i* for each seed. We then estimated 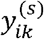 and 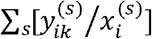, which shows the concordance in the assignation of each individual to a cluster across the different seeds. To summarize this information, we analyzed all the individual sums, for each seed and each cluster *k* (Fig. S7A and C), as well as the number of individuals who were not assigned to the same cluster, between one seed and all the other seeds (Fig. S7B and D).

Based on these analyses, we chose the seed that showed the highest consensus among seeds (Fig. S7) and in which the largest number of samples are robustly assigned (Fig. S6) (seed number 289 for Analysis 1 and 161 for Analysis 2). Then, we defined the maximum *k* value based on the number of individuals who are robustly assigned to a cluster for each *k* value and we focused our analyses on this value. Finally, once we chose a seed and a *k* value, we removed samples that showed some uncertainty in the cluster assignation (*P_i_* ≤ 0.9). Following this process, we observed that the robustness decreases from *K*_FS_ = 20 and we removed 3 samples from Analysis 1 (Vanuatu). For the reference dataset (Analysis 2), all samples showed a *P_i_* > 0.9 across all *k* values, therefore we used genetic clusters from the *k* value that best suited the hypotheses to test (*K*_FS_ = 25) (Table S5).

For the subsequent analyses, we redefined populations based on fineSTRUCTURE results and therefore on the genetic data itself. This approach is rather trivial when applied to continental populations, like the reference populations presented here, because fineSTRUCTURE clusters coincide broadly with the geographical or cultural categories assigned to the populations. Yet, this strategy has proven useful when complex admixture shapes the genetic structure [59, 60], or when the studied populations are sampled from a small geographic region [20], in which case, defining populations based on geography or culture may lead to a biased view of the genetic structure of the populations.

#### Ancestry estimation

We estimated admixture proportions for each ni-Vanuatu individual using SOURCEFIND [61]. We first ran ChromoPainter using the “donors” mode: we considered each ni-Vanuatu individual as a recipient only and defined donor populations by using the genetic clusters defined above (Analysis 2). We ran SOURCEFIND by considering all surrogates (“all surrogates” analysis; Fig. S15 and Table S5) or by limiting as much as possible the number of surrogates (“limited surrogates” analysis). In the latter case, we considered as surrogates West_Eurasia1, West_Eurasia2, West_Eurasia3, EastAsia1, EastAsia2, EastAsia3, Atayal, Paiwan, Southeast_Asia, Cebuano, RenBell, Tikopia, PNG1, PNG2 and PNG_SGDP. Then, we summed the estimated ancestry proportions for these surrogates and grouped them as follows: Taiwanese.Philippines (Atayal, Paiwan, Cebuano), EastAsian_mainland (EastAsia1, EastAsia2, EastAsia3, Southeast_Asia), Polynesian (RenBell, Tikopia), Papuan-related (PNG1, PNG2 and PNG_SGDP) and European (West_Eurasia1, West_Eurasia2, West_Eurasia3). The East Asian-related ancestry was estimated as the sum of Taiwanese.Philippines group and Polynesian group (the EastAsian_mainland was negligible, with a maximum proportion = 0.007). To facilitate visualization, we excluded an individual born in Futuna and living in Malekula, who shows a Polynesian ancestry of 0.745 in figures showing ancestry proportions (Fig. 2, Fig. S8-11). The next individual carrying the highest Polynesian ancestry shows 0.383. For both the “all surrogates” and “limited surrogates” analyses, we ran 200,000 MCMC iterations with a burn-in of 50,000 iterations, sampling every 5,000 iterations and not allowing for self-copy. We estimated the final ancestry of each ni-Vanuatu sample as the mean of the 30 sampled MCMC runs.

#### Admixture date estimation

We estimated admixture dates using GLOBETROTTER [62]. For this analysis, we ran ChromoPainter using the “donors” mode, by considering the ni-Vanuatu genetic clusters defined in (Analysis 1) as recipients and all the reference populations (Analysis 2) as both recipients and donors. Surrogates were the same as those considered for the “limited surrogates” SOURCEFIND analysis. For each of the 20 recipient ni-Vanuatu clusters, we performed 100 bootstraps, which are implemented as resamplings of chromosomes among the available samples. We set the “Null.ind” option to 1. We assumed a generation time of 28 years, to estimate dates from the number of generations. We also dated admixture in each ni-Vanuatu sample following the same approach.

#### Sex-biased admixture

We studied sex-biased admixture by comparing ancestry proportions estimated for the autosomes and the X chromosome. We estimated ancestry proportions using RFMix v.1.5.4 [33], considering as ancestral populations New Guinean highlanders (approximating Papuan-related ancestry) and Taiwanese Indigenous peoples and the Cebuano from the Philippines (approximating East Asian-related ancestry). We ran the “TrioPhased” algorithm, allowing for phase correction and 3 EM iterations. The window size was set at 0.03 cM [18] and the admixture date was set at 50 generations. The same parameters were used for the X chromosome and the autosomes. We combined the X chromosome haploid data of males from the same island to obtain diploid individuals. We filtered out SNPs with an RFMix posterior probability < 0.9, those within centromeres and within 2 Mb from the telomeres. We then estimated East Asian-related and Papuan-related ancestry in the autosomes and the X chromosome, separately. We estimated *α_f_* and *α_m_*, that is, the proportion of female and male ancestors of ni-Vanuatu who carried Papuan-related ancestry, respectively, by assuming that admixture proportions have reached equilibrium values [34]. In such a case, *α_f_* = 3*α_x_* – 2*α_auto_* and *α_m_* = 4*α_auto_* – 3*α_X_*, where *α_auto_* and *α_X_* are the average Papuan-related ancestry proportions estimated on the autosomes and the X chromosome, respectively.

To avoid potential biases due to SNP ascertainment, we also studied sex-biased admixture by comparing the autosomes and the sex chromosomes of 179 ni-Vanuatu sequenced at 30× coverage [18]. Because genotypes on the X chromosome were not called in the previous study, we mapped *fastq* files on the X chromosome of the human reference genome (version hs37d5) and performed genotype calling for X-linked variants, as previously described [18], setting the ploidy parameter to 1 for males and 2 for females. We kept only biallelic SNPs and filtered *vcf* files following GATK best practices [63]: QualByDepth < 2; FisherStrand value > 60; StrandOddsRatio > 3; RMSMappingQuality < 40; MappingQualityRankSumTest < −12.5; and ReadPosRankSum < −8. We also removed genotypes with a DP < 10 for females and < 3 for males, and those with a GQ < 30. We removed PAR1, PAR2 and XTR regions according to the annotations provided by the UCSC browser, and amplicon regions as reported in [64]. We filtered out SNPs that were not in Hardy-Weinberg equilibrium in females (*P*-value < 0.0001), those with a missingness > 0.05 and those with a MAF < 0.01. We estimated mtDNA haplogroups with MToolBox [65] from the *bam* and *fastq* files, and Y chromosome haplogroups were estimated with Yleaf [66].

#### Inferring migrations within Vanuatu

We used the genetic structure inferred by fineSTRUCTURE to study migration patterns among Vanuatu islands. First, we assigned each genetic cluster to one or more islands, each time more than 25% of the sampled individuals inhabiting the island were assigned to that cluster. We then identified “outlier” individuals who inhabit an island but are assigned to a genetic cluster that is prevalent in another island. These individuals, either themselves or their close ancestors, likely migrated from the island/s where their genetic cluster is predominant to their current place of residence. We removed from these analyses Vanuatu_8 and Vanuatu_9 clusters (*K*_FS_ = 20) because those clusters are driven by recent European gene flow, as well as Vanuatu_15 because no island could be assigned to this cluster when using the 25% rule. To study sex-specific migrations, we calculated the proportion of female (male) migrants by dividing the number of “outlier” females (males) by the number of females (males) in each cluster. We did not include clusters where the number of males or females was < 2.

#### Tests for exogamy

We tested if spouses show lower genetic relatedness than non-spouses by testing if the average kinship coefficient between 287 observed couples is higher or lower than expected by chance, using either permutations or a logistic regression model. Kinship coefficients were computed between all possible pairs of individuals using KING v.2.1 (Table S7). When using permutations, we accounted for isolation by distance by sampling random pairs of males and females among individuals born in the same island (Fig. S24A) or living in the same village (Fig. S24B). Because marriages between first-degree related individuals are very unlikely (they are indeed not observed in the data), we excluded from this analysis individuals who are first-degree relatives, but also tested how the inclusion of these individuals affect the results (Fig. S24C). We performed 10,000 permutations for each island. In each permutation, we sampled equally many random pairs as there were observed pairs in the island. We calculated *P*-values by comparing the average kinship coefficient among the observed couples to the null distribution. The null distribution was made of 10,000 average kinship coefficients between randomly sampled pairs of individuals from the same island or village.

Based on the sample scheme described above, we designed a logistic regression model that could be generalized to multiple explanatory variables, such as ancestry. Let the variable *i* index all possible pairs of males and females in the dataset, except pairs of individuals who are first-degree relatives, and define the dependent binary variable *y_i_* with *Y_i_* = 1 if *i* indexes an observed pair of spouses and *Y_i_* = 0 otherwise. Define the probability that a pair is an observed couple *p_i_*(*x*) = *P*(*Y_i_* = 1), and introduce *ϕ_i_* as the kinship coefficient between individuals in the *i*:th pair. We estimated the effect of kinship on mate choice by the parameter β^ϕ^ in the logistic regression model

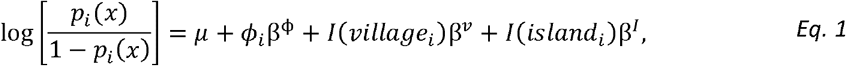

where *I*(*village_i_*) = 1 if individuals of pair *i* are from the same village and *I*(*village_i_*) = 0 otherwise, and *I*(*island_i_*) = 1 if individuals of pair *i* are from the same island and *I*(*island_i_*) = 0 otherwise. As a sensitivity analysis, we also considered a model in which pairs of first-degree related individuals were included (Fig. S25).

#### Tests for ancestry-based assortative mating

We extended the logistic regression model shown in *Eq. 1* to test for ancestry-based assortative mating, by testing if the genetic ancestry of spouses is more similar than that of non-spouses, accounting for population structure and relatedness avoidance. Let 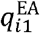 and 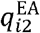 be the proportion of East Asian-related ancestry for the individuals 1 and 2 of pair *i* and introduce the variable 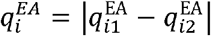. We estimated the effect of having similar proportions of East Asian-related ancestry between two individuals on the probability that a pair is an observed couple *p_i_*(*x*) by the parameter β^*q*^ in the logistic regression model,

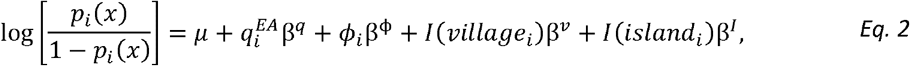

where we use the same notations as in *Eq. 1*. A negative (positive) effect size β^q^ is interpreted as evidence for assortative (disassortative) mating according to ancestry. Effect sizes for other ancestries were calculated similarly. The same model was also tested in ni-Vanuatu originating or not from islands where Polynesian languages are spoken. Note that in standard regression analyses, such as those used in GWAS, population stratification is usually corrected by principal components of the genetic relatedness matrix; here we have taken a more general approach by decomposing the genetic structure in two variables: the kinship and the ancestry, and have studied how both variables affect mate choice independently.

#### Tests for SNP-based assortative mating

We extended the logistic regression model shown in *Eq. 1* to test for SNP-based assortative mating, by testing if the genotypes at a given SNP are more similar between spouses than non-spouses, accounting for population structure, ancestry-based associated mating and relatedness avoidance. Define, for each SNP *s* and each pair *i* of individuals, the allele sharing distance (ASD) *d_i,s_* as *d_i,s_* = 0 if both alleles are identical between individuals, *d_i,s_* = 1 if only one allele is identical and *d_i,s_* = 2 if none of the alleles are identical. We estimated the association between SNP *s* and mate choice by the parameter 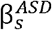 in the model

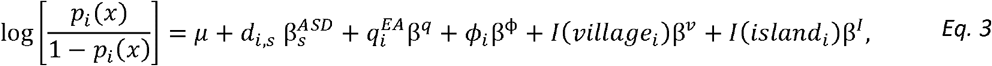

where we use the same notations as in *Eq. 2*. A negative (positive) effect size 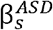 indicates higher (lower) similarity at SNP *S* between the members of the observed couples than between non-couples, in line with SNP-based assortative (disassortative) mating.

#### Tests for trait-based assortative mating

We tested for trait-based assortative mating by testing if genotypes of spouses are more similar or dissimilar at trait-associated SNPs, relative to non-associated SNPs. We obtained GWAS summary statistics for 8 candidate traits from the UK Biobank database (http://www.nealelab.is/uk-biobank). Candidate traits include traits relating to morphology and physical appearance. Let 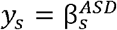, where 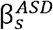 is the effect size of SNP *s* on mate choice estimated by *Eq. 1*. We estimated if trait-associated SNPs are more similar or dissimilar between spouses by the parameter β^*trait*^ in the model

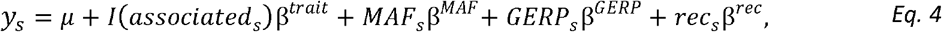

where *I*(*associated_s_*) = 1 if SNP *s* is significantly associated with the candidate trait in GWAS (with GWAS *P*-value < 5×10^-8^) and *I*(*associated_s_*) = 0 otherwise, *MAF_s_* is the minor allele frequency of SNP *s* in ni-Vanuatu, *GERP_s_* is the Genomic Evolutionary Rate Profiling (GERP) score of SNP *s* and *rec_s_* is the interpolated recombination rate between SNP *s* and SNP *s* – 1, estimated in cM/Mb from the 1000 Genomes Phase 3 combined recombination map. We considered that there is trait-based assortative (disassortative) mating if *β^trait^* is significantly negative (positive). We adjusted *P*-values with the Bonferroni correction for multiple testing, to account for the number of traits tested.

## ACKNOWLEDGMENTS

We thank all volunteers in Vanuatu who kindly provided samples and participated in this research. We thank the Ministry of Health of the Republic of Vanuatu (especially Myriam Abel, Rose Bahor, Yvannah Taga, Maturine Tary and the former Minister Jerôme Ludvaune), the Vanuatu Family Health Association (especially Blandine Boulekone and Marie Nickllum Wala), the WHO Office in Port Vila (especially Corinne Capuano) and the Institut Pasteur of New Caledonia at Nouméa (especially Paul M. V. Martin and Eliane Chungue), for their continuous support and interest in this work. We are particularly thankful to Helene Walter, Woreka Mera, Françoise Charavay and Sonia Treptow for their assistance in collecting samples. We thank S. Créno and the HPC Core Facility of Institut Pasteur (Paris) for the management of computational resources. L.R.A. was funded by a Pasteur-Roux-Cantarini fellowship from the Institut Pasteur. The laboratory of Human Evolutionary Genetics is supported by the Institut Pasteur, the Collège de France, the CNRS, the Fondation Allianz-Institut de France, the French Government’s Investissement d’Avenir programme, Laboratoires d’Excellence ‘Integrative Biology of Emerging Infectious Diseases’ (ANR-10-LABX-62-IBEID) and ‘Milieu Intérieur’ (ANR-10-LABX-69-01), the Fondation de France (n°00106080), and the Fondation pour la Recherche Médicale (Equipe FRM DEQ20180339214) and the French National Research Agency (ANR-19-CE35-0005).

## AUTHOR CONTRIBUTIONS

Conceptualization: L.R.A., E.P.; L.Q.-M.; Methodology: L.R.A., J.B., A.M.S., E.P.; Software: L.R.A., M.R.; Formal Analysis: L.R.A.; Data Generation: C.H., L.L.; Resources: O.C., A.G.; Writing – Original Draft: L.R.A., L.Q.M., E.P.; Writing – Review & Editing: L.R.A., J.B., J.C., J.M.R., M.R., A.M.S., H.C., A.F., F.V., L.Q.-M., E.P.; Visualization: L.R.A.; Supervision: L.Q.-M., E.P.; Funding Acquisition: L.Q.M.

## SUPPLEMENTAL FIGURE LEGENDS

**Figure S1. Genetic structure of ni-Vanuatu and a selection of worldwide populations.**

(A) ADMIXTURE analyses of the SNP array data for 1,439 ni-Vanuatu, together with whole genome sequences of other populations from the Pacific region [18]. (B) ADMIXTURE cross-validation errors for different *K*_ADM_ values and 10 iterations with different seeds, indicated by colors.

**Figure S2. Principal Component Analyses of the SNP array dataset for ni-Vanuatu.** Each point is an individual, colored according to the geographical latitude coordinates of their island of residence, as in Fig. 1A. Percentages indicate the proportion of variance explained. PC1 and PC2 are shown in Fig. 1. Black crosses and points indicate projected ancient samples from Vanuatu dated to the Lapita and Post-Lapita periods, respectively.

**Figure S3. *F*_ST_ matrix between ni-Vanuatu and other worldwide populations.** *F*_ST_ values and their significance were obtained with the StAMPP R package [58]. The lower triangle of the matrix indicates *F*_ST_ values, whereas the upper triangle indicates *P*-values. All values were significant (*P*-value < 0.01, using 100 bootstraps).

**Figure S4. Map showing the clustering of ni-Vanuatu into *K_FS_* = 2 to 18 genetic clusters, according to fineSTRUCTURE.** Each point indicates an individual, located according to their village of residence. Colors indicate clusters, so that the closer the colors, the closer the clusters. Noise was added to sampling locations to facilitate visualization.

**Figure S5. Sample sizes of each genetic cluster, according to fineSTRUCTURE at *K*_FS_ = 20.** Filled bars indicate the number of individuals who live in a village where Polynesian languages are spoken.

**Figure S6. Robustness of fineSTRUCTURE analyses.** (A) Number of individuals below a 0.9 threshold of confidence in the assignation to a specific cluster (y-axis), as a function of a number of clusters *K*_FS_ that varies between 1 and 125 (x-axis). Colors indicate the 5 different seeds used for the MCMC algorithm. (B) Zoom-in of the plot in (A), for *K*_FS_ between 1 and 50. As expected, the robustness on the assignation of each individual to a genetic cluster decreases with increasing *K*_FS_ values (i.e., more samples are below the 0.9 threshold). (C) Robustness as measured by the average of Σ_m_(*Y*_i_ / *X*_i_) (Methods), as a function of a number of clusters *K* that varies between 1 and 125 (x-axis). (D) Zoom-in of the plot in (C), for *K*_FS_ between 1 and 50. The shaded areas indicate the standard deviation of ∑_m_(*Y*_i_ / *X*_i_) for each *K*_FS_ value.

**Figure S7. Robustness of fineSTRUCTURE analyses across seeds, for the chosen *K*_FS_ values.** (A,C) Heatmaps showing the average of Σ_S_(*Y*_i_ / *X*_i_) across clusters for (A) the ni-Vanuatu (Analysis 1) (*K*_FS_ = 20) and (C) the reference populations (Analysis 2) (*K*_FS_ = 25). Each row corresponds to one seed (the number of which is circled in black), and each column corresponds to a genetic cluster. Colors indicate the mean Σ_S_(*Y*_i_ / *X*_i_) for each cluster and seed. (B,D) Heatmaps showing the mismatch between each seed and the four other seeds for (B) the ni-Vanuatu (Analysis 1) (*K*_FS_ = 20) and (D) the reference populations (Analysis 2) (*K*_FS_ = 25). Each row corresponds to the seed used as reference to estimate ∑_S_(*Y*_i_ / *X*_i_). Each column corresponds to a genetic cluster. Colors indicate the proportion of individuals who are assigned to a different cluster in the other seeds.

**Figure S8. Ancestry proportions in ni-Vanuatu, estimated by SOURCEFIND.** Each point indicates an individual, colored according to their ancestry proportions. The East Asian-related ancestry is the sum of the Taiwan. Philippines and Polynesian ancestry. Noise was added to sampling locations to facilitate visualization.

**Figure S9. Correlations between East Asian-related and Polynesian ancestry inferred by SOURCEFIND.** (A) Each individual is colored according to their island of residence. (B) The black points indicate the individuals living in villages where Polynesian languages are spoken, the rest of the samples are colored in gray. The blue line indicates the regression line considering all the individuals from Vanuatu. The gray area indicates the 95% confidence level.

**Figure S10. Distributions of ancestry proportions in ni-Vanuatu, estimated by SOURCEFIND, for each island and ancestry.** The line, box, whiskers and points, respectively, indicate the median, interquartile range (IQR), 1.5*IQR and outliers. Colors indicate the geographical latitude coordinates of each island, as in Fig. 1A.

**Figure S11. Distributions of ancestry proportions of ni-Vanuatu among genetic clusters, according to fineSTRUCTURE at *K*_FS_ = 4.** The line, box and whiskers, respectively, indicate the median, interquartile range (IQR) and 1.5*IQR. We tested significant differences in ancestry between each cluster using a Wilcoxon test with Bonferroni correction. \**P* < 0.05, \*\**P* < 0.01, \*\*\**P* < 0.001, \*\*\*\**P* < 0.0001, ns: *P* > 0.05.

**Figure S12. Map showing admixture dates estimated per individual with GLOBETROTTER.**

**Figure S13. Allele sharing of ni-Vanuatu with Near Oceanians, according to *f*_4_-statistics.** The x-axis indicates *f*_4_(X, New Guinean Highlanders; EastAsian, Australian), which measures allele sharing of ni-Vanuatu and Near Oceanians (X) with East Asians. The y-axis indicates *f*_4_(X, New Guinean Highlanders; Near Oceanian, EastAsian), which measures allele sharing of ni-Vanuatu and Near Oceanians (X) to each Near Oceanian population tested (Nakanai, Nasioi, Baining), relative to their allele sharing with New Guinean Highlanders. The regression line is estimated considering only modern individuals from Vanuatu. The bars show two standard errors. Deviations upper from the regression line indicate a higher affinity with the Near Oceanian population tested.

**Figure S14. Principal Component Analysis (PCA) of ni-Vanuatu, together with other populations from Near Oceania.** The modern and ancient samples from Vanuatu were projected into a PCA of a previous SNP array data set [41].

**Figure S15. Ancestry proportions in ni-Vanuatu, estimated by SOURCEFIND, considering all the reference populations as possible surrogates (Table S4).** The bar plot indicates the mean value across individuals, for each ancestry in each island of residence.

**Figure S16. Allele sharing of ni-Vanuatu with Polynesians, according to *f*_4_-statistics.** The x-axis indicates *f*4(X, New Guinean Highlanders; EastAsia, Australian), which measures allele sharing of ni-Vanuatu and Near Oceanians (X) with East Asians. The y-axis indicates *f*_4_(X, EastAsian; Polynesian, Tolai), which measures allele sharing of ni-Vanuatu and Near Oceanians (X) with Polynesians (Rennell and Bellona [RenBel], Tikopia, Tonga), relative to their East Asian ancestry. The regression line is estimated considering only modern individuals from Vanuatu. The bars show two standard errors. Deviations upper from the regression line indicate a higher affinity with the Polynesian population tested.

**Figure S17. Sex-biased admixture in Vanuatu.** (A) X-to-autosome ratio of Papuan-related ancestry proportions in ni-Vanuatu who speak non-Polynesian or Polynesian languages, estimated using the SNP array data (Wilcoxon test *P*-value < 1.36×10^-5^, after Bonferroni correction). (B) Papuan-related ancestry proportions in ni-Vanuatu, estimated for the 22 autosomes and the X chromosome separately, using the high-coverage genome sequencing data obtained for a subset of 179 ni-Vanuatu [18].

**Figure S18. Self-reported migrations among ni-Vanuatu spouses.** (A) Map of self-reported migrations for females (blue) and males (green), separately. The arrows connect the place of birth of individuals to their place of residence. (B) Barplot showing the number of females (blue) and males (green) among ni-Vanuatu spouses who reported to migrate during their lifetime.

**Figure S19. Proportions of female and male ni-Vanuatu migrants, based on genetic clusters or self-reported information.**

**Figure S20. Comparison of kinship levels among ni-Vanuatu living in the same village, the same island or the entire archipelago.** Significance of the differences was tested by a Wilcoxon test.

**Figure S21. Genetic relatedness and islands of residence of related ni-Vanuatu.** Each line connects two individuals who are related up to third degree. Islands are order from north to south (clockwise).

**Figure S22**. **Degrees of genetic relatedness of ni-Vanuatu from the different Vanuatu islands.** Bar plots indicate the proportion of pairs of related individuals according to the inferred degree of genetic relatedness, in each island. The black points and indicate the mean kinship per island. Significance was tested by a Fisher’s exact test. \**P* < 0.05, \*\**P* < 0.01, \*\*\**P* < 0.001.

**Figure S23. Genetic relatedness and islands of residence of ni-Vanuatu spouses.** Each line connects the two spouses, and the color indicates the degree of genetic relatedness. Islands are order from north to south (clockwise).

**Figure S24**. **Kinship among spouses and random pairs of individuals.** Kinship coefficients among spouses and random pairs of individuals from the same (A) island (excluding first-degree related pairs), (B) village (excluding first-degree related pairs) and (C) village (including first-degree related pairs). The orange vertical line indicates the average kinship coefficient among spouses. The blue density curve indicates the null distribution of kinship coefficients estimated from the sampling of random pairs of individuals. Each data point of the null distribution is estimated as the mean kinship coefficient in randomly sampled pairs of individuals. Of note, negative kinship coefficients are expected when the two compared individuals are from different populations.

**Figure S25**. **Effects of geographical location and kinship on mate choice.**. (A) Effect sizes estimated when not including the village of residence as a predictor. (B) Effect sizes estimated when including the village of residence as a predictor and excluding first-degree related pairs from the analysis. (C) Effect sizes estimated when including the village of residence as a predictor and including first-degree related pairs from the analysis. (A-C) Effect sizes of the logistic regression modelling the probability of mating as a function of three predictors: the island of birth, the village of residence and kinship coefficients.

**Figure S26. Q-Q plots of observed and expected *P*-values of a regression model testing for SNP-based assortative mating**. (A) *P*-values are from a model that does not control for population structure. (B) *P*-values are from a model that controls for population structure (island of birth, kinship and ancestry). The shaded area indicates the 95% confidence interval.

