## Supplementary Information for "The genomic landscape of contemporary western Remote Oceanians"

**Supplemental Information for**
### The genomic landscape of contemporary western Remote Oceanians

Lara R. Arauna, Jacob Bergstedt, Jérémy Choin, Javier Mendoza-Revilla, Christine Harmant, Maguelonne Roux, Alex Mas-Sandoval, Laure Lémée, Heidi Colleran, Alexandre François, Frédérique Valentin, Olivier Cassar, Antoine Gessain, Lluis Quintana-Murci, Etienne Patin

**This PDF file includes:**

Figures S1 to S26

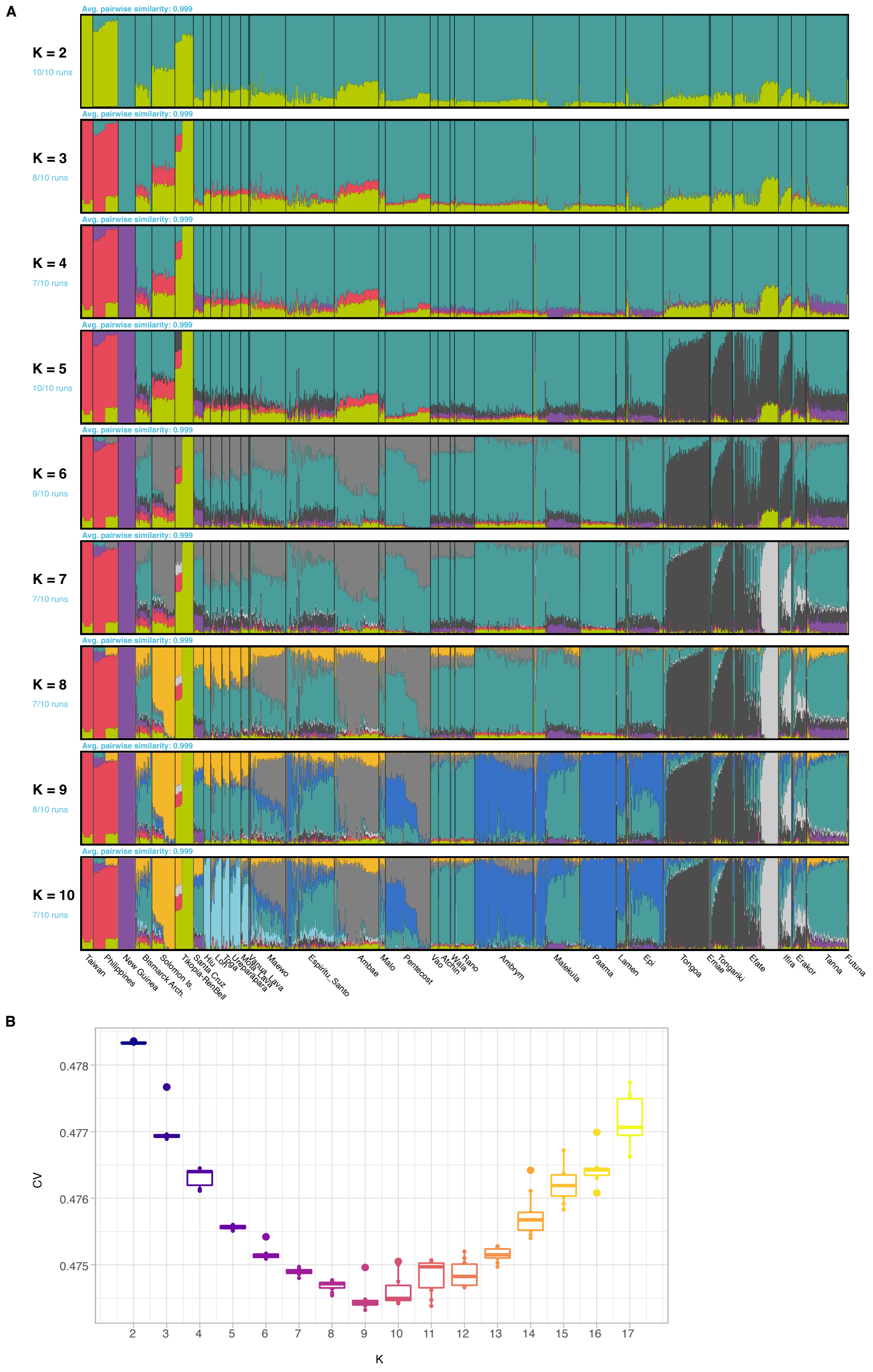


**Figure S1. Genetic structure of ni-Vanuatu and a selection of worldwide populations.**
(A) ADMIXTURE analyses of the SNP array data for 1,439 ni-Vanuatu, together with whole genome sequences of other populations from the Pacific region [18].
(B) ADMIXTURE cross-validation errors for different $K_{ADM}$ values and 10 iterations with different seeds.

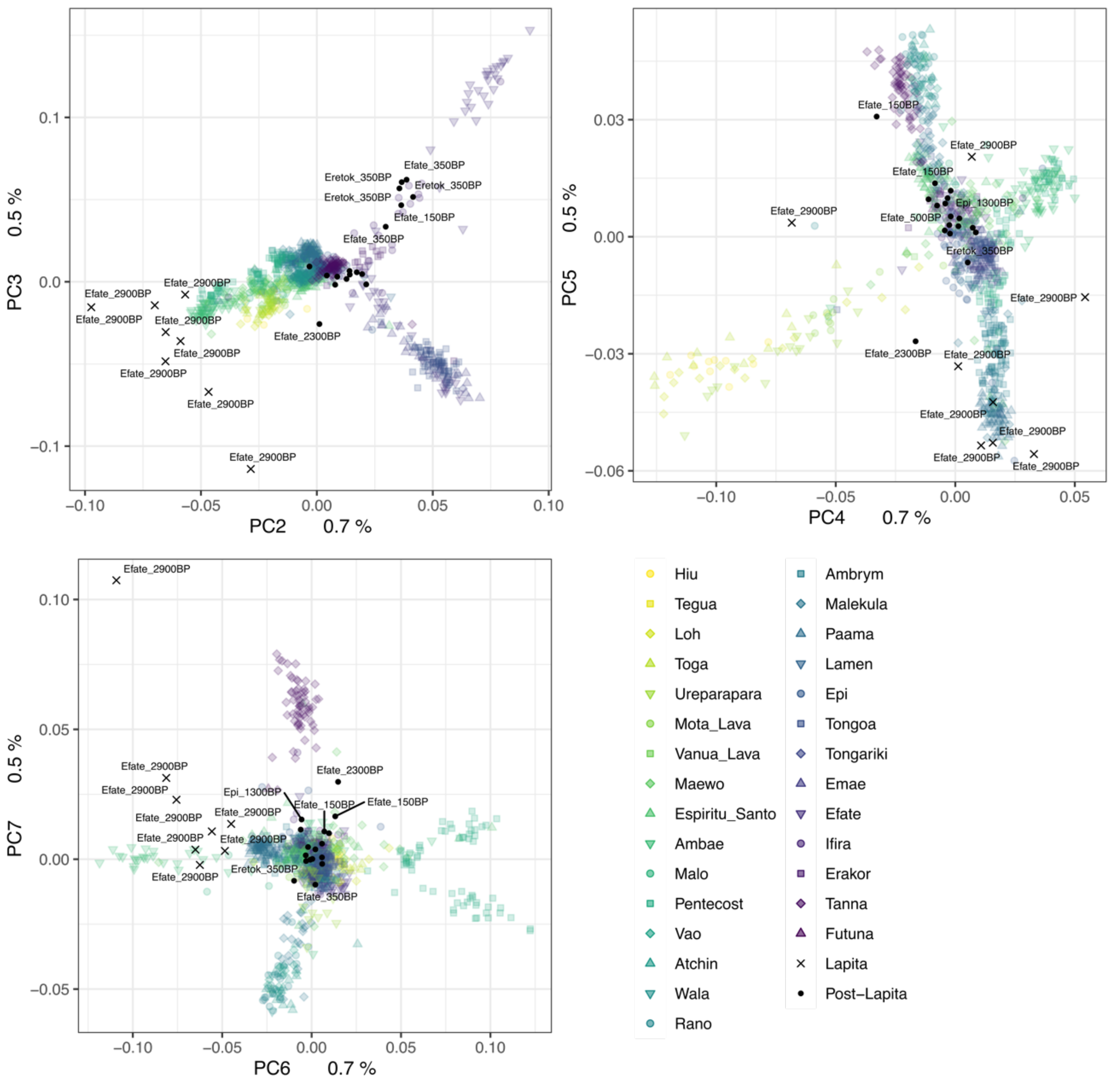


**Figure S2.** Principal Component Analyses of the SNP array dataset for ni-Vanuatu. Each point is an individual, colored according to the geographical latitude coordinates of their island of residence, as in Fig. 1A. Percentages indicate the proportion of variance explained. PC1 and PC2 are shown in Fig. 1. Black crosses and points indicate projected ancient samples from Vanuatu dated to the Lapita and Post-Lapita periods, respectively.

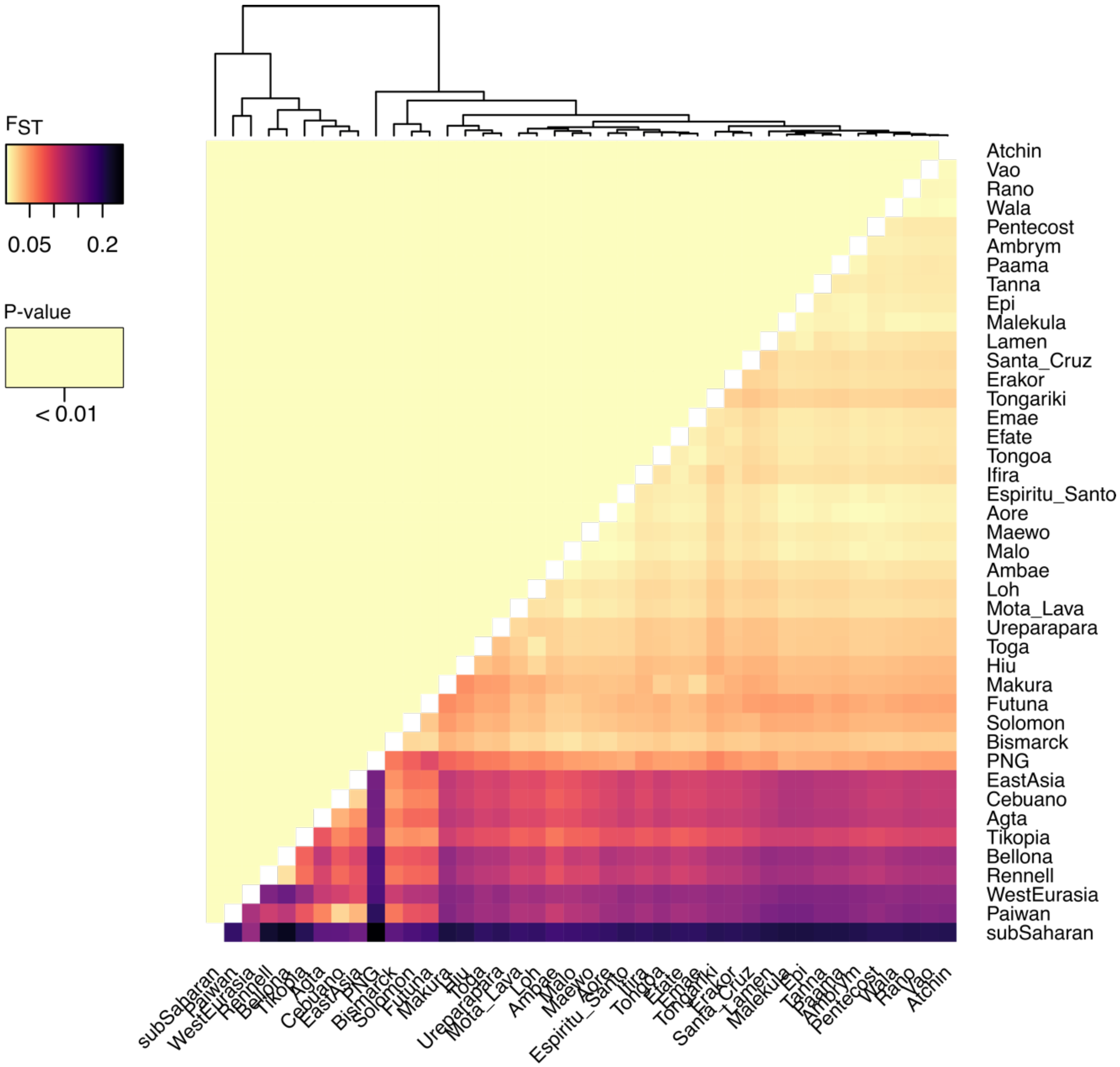


**Figure S3. $F_{ST}$ matrix between ni-Vanuatu and other worldwide populations.**
$F_{ST}$ values and their significance were obtained with the StAMPP R package [58]. The lower triangle of the matrix indicates $F_{ST}$ values, whereas the upper triangle indicates $P$-values. All values were significant ($P$-value < 0.01, using 100 bootstraps).

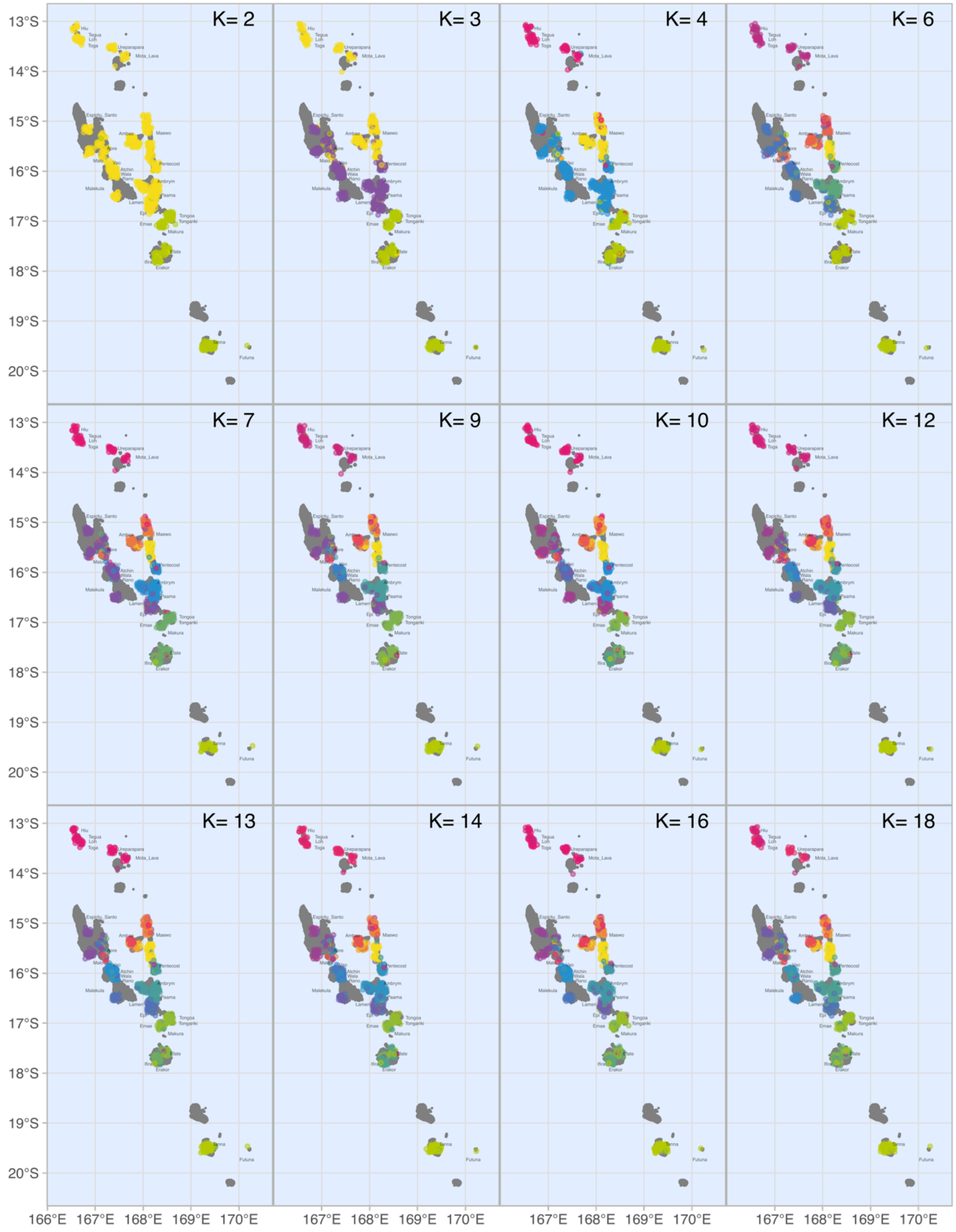


**Figure S4. Map showing the clustering of ni-Vanuatu into $K_{FS}$ = 2 to 18 genetic clusters, according to fineSTRUCTURE.**

Each point indicates an individual, located according to their village of residence. Colors indicate clusters, so that the closer the colors, the closer the clusters. Noise was added to sampling locations to facilitate visualization.

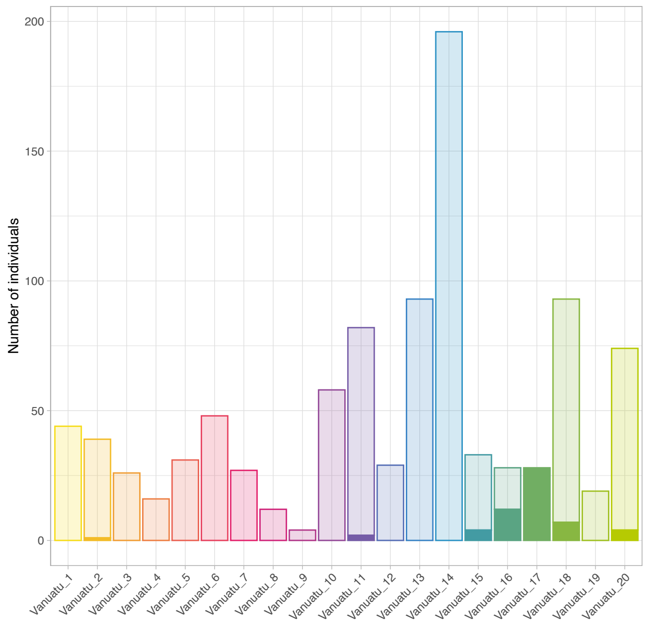


**Figure S5. Sample sizes of each genetic cluster, according to fineSTRUCTURE at $K_{FS}$ = 20.** Filled bars indicate the number of individuals who live in a village where Polynesian languages are spoken.

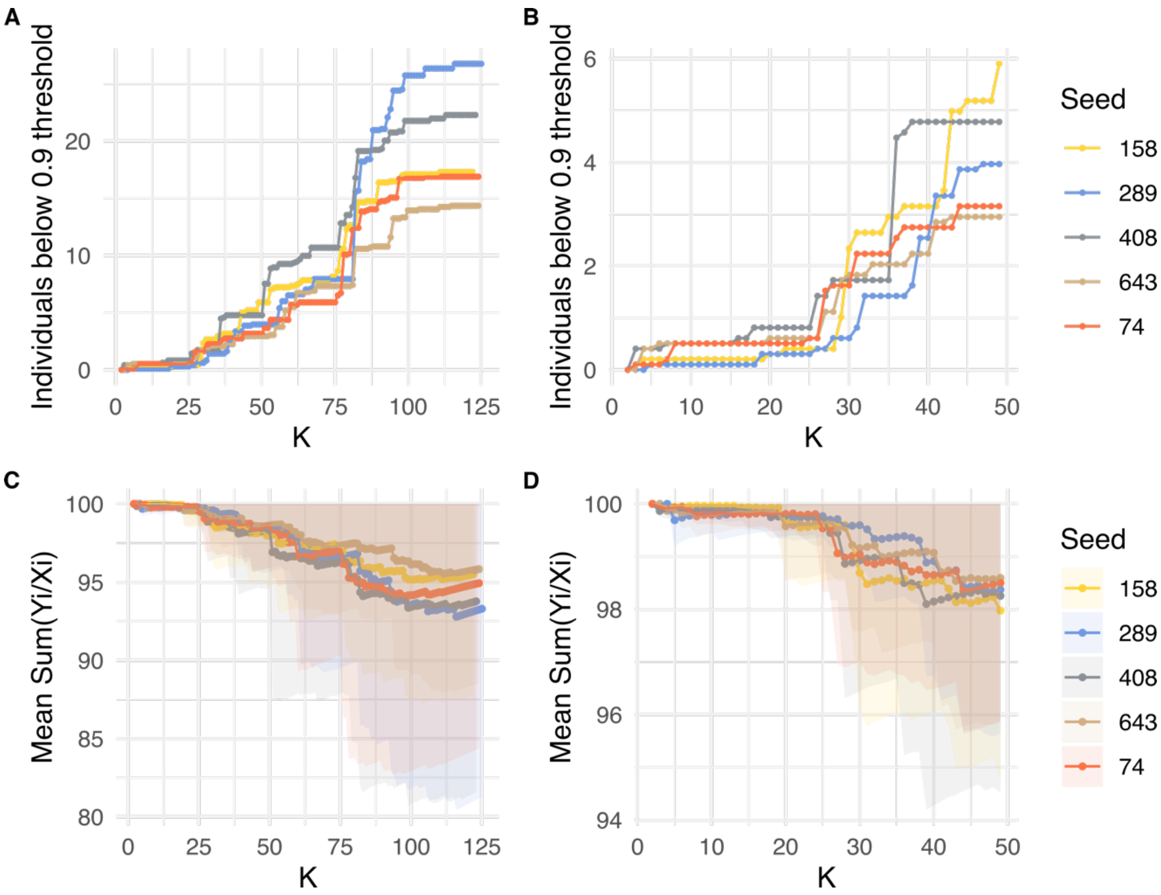


**Figure S6. Robustness of fineSTRUCTURE analyses.**
(A) Number of individuals below a 0.9 threshold of confidence in the assignation to a specific cluster (y-axis), as a function of a number of clusters $K_{FS}$ that varies between 1 and 125 (x-axis). Colors indicate the 5 different seeds used for the MCMC algorithm.
(B) Zoom-in of the plot in (A), for $K_{FS}$ between 1 and 50. As expected, the robustness on the assignation of each individual to a genetic cluster decreases with increasing $K_{FS}$ values (i.e., more samples are below the 0.9 threshold).
(C) Robustness as measured by the average of $\Sigma_m(Y_i / X_i)$ (Methods), as a function of a number of clusters $K$ that varies between 1 and 125 (x-axis).
(D) Zoom-in of the plot in (C), for $K_{FS}$ between 1 and 50. The shaded areas indicate the standard deviation of $\Sigma_m(Y_i / X_i)$ for each $K_{FS}$ value.

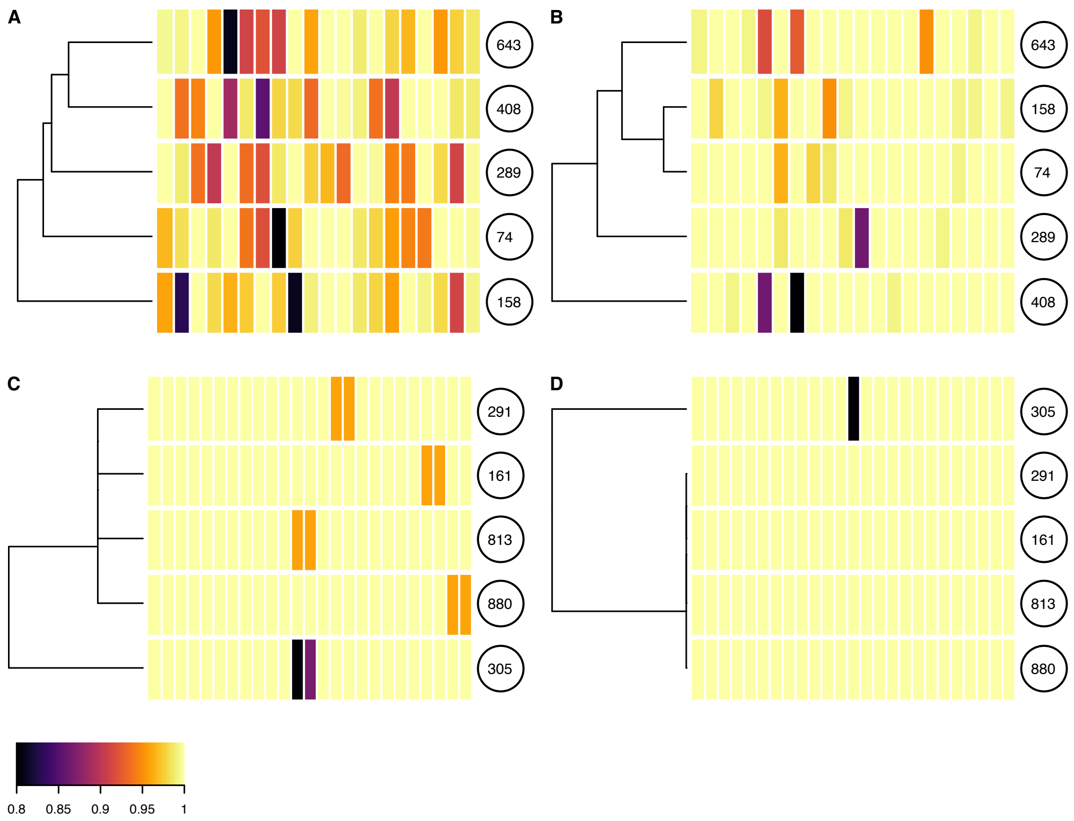


**Figure S7. Robustness of fineSTRUCTURE analyses across seeds, for the chosen $K_{FS}$ values.**

(A,C) Heatmaps showing the average of $\Sigma_S(Y_i / X_i)$ across clusters for (A) the ni-Vanuatu (Analysis 1) ($K_{FS}$ = 20) and (C) the reference populations (Analysis 2) ($K_{FS}$ = 25). Each row corresponds to one seed (the number of which is circled in black), and each column corresponds to a genetic cluster. Colors indicate the mean $\Sigma_S(Y_i / X_i)$ for each cluster and seed.

(B,D) Heatmaps showing the mismatch between each seed and the four other seeds for (B) the ni-Vanuatu (Analysis 1) ($K$ = 20) and (D) the reference populations (Analysis 2) ($K_{FS}$ = 25). Each row corresponds to the seed used as reference to estimate $\Sigma_S(Y_i / X_i)$. Each column corresponds to a genetic cluster. Colors indicate the proportion of individuals who are assigned to a different cluster in the other seeds.

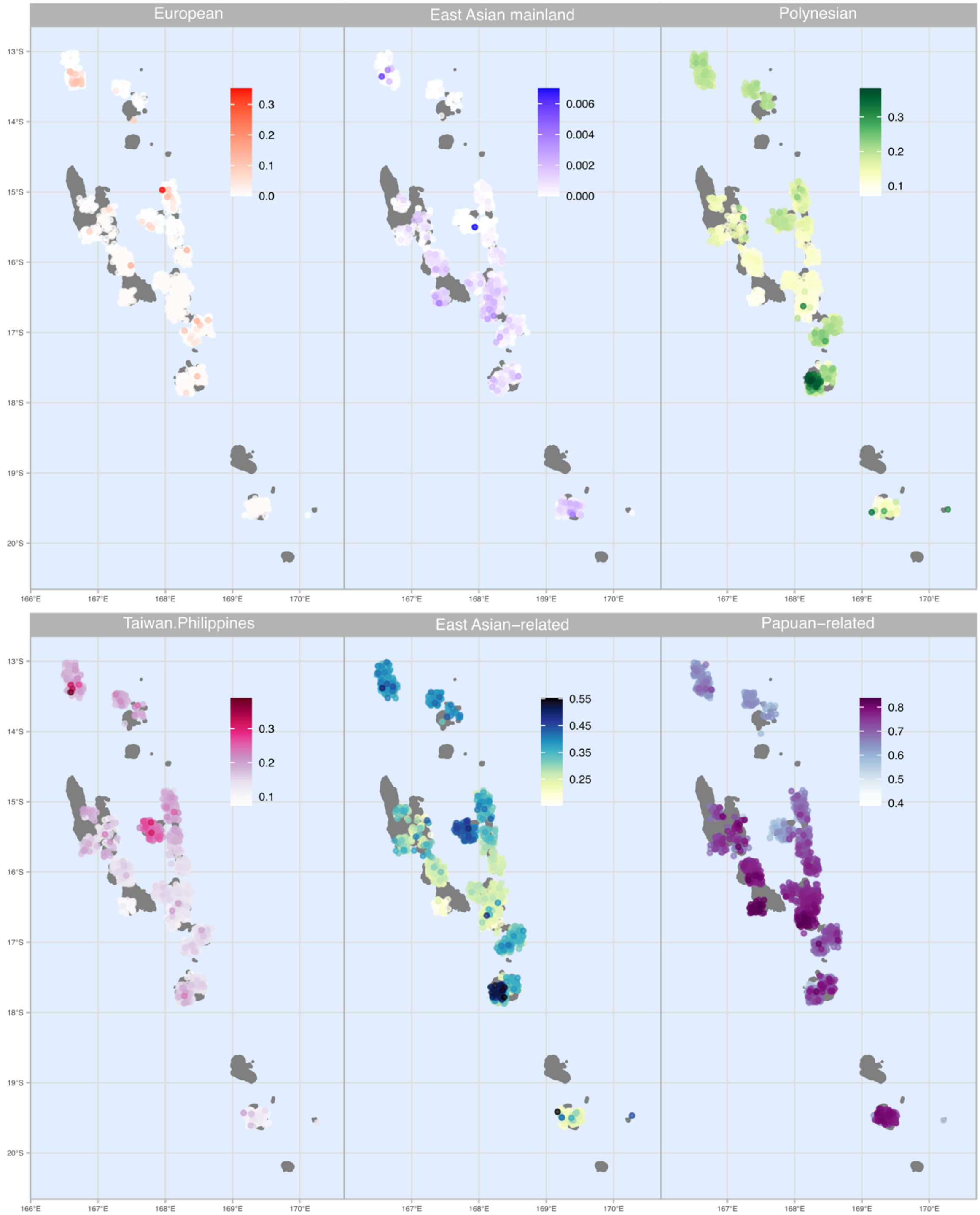


**Figure S8. Ancestry proportions in ni-Vanuatu, estimated by SOURCEFIND.**
Each point indicates an individual, colored according to their ancestry proportions. The East Asian-related ancestry is the sum of the Taiwan.Philippines and Polynesian ancestry. Noise was added to sampling locations to facilitate visualization.

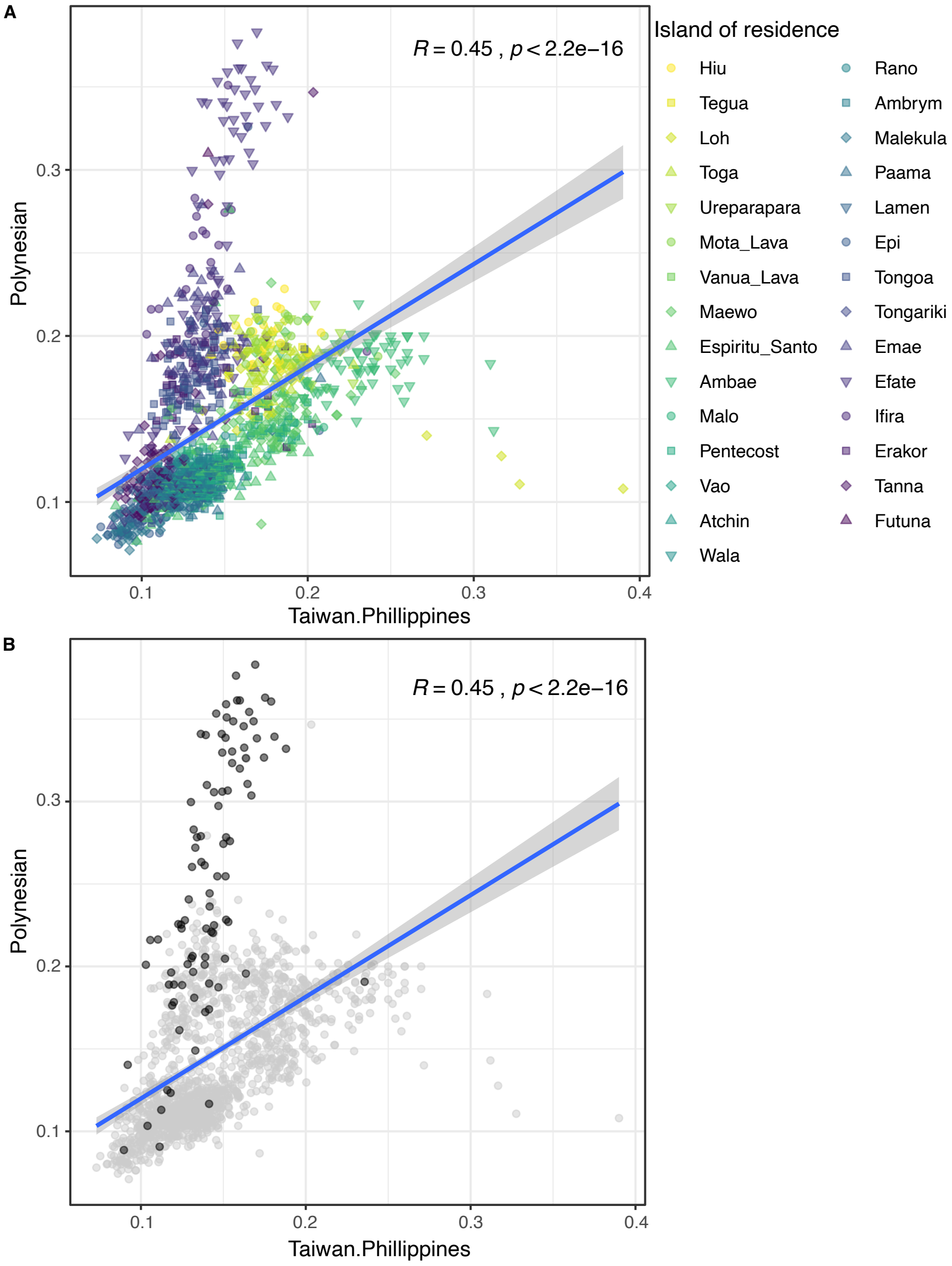


**Figure S9. Correlations between East Asian-related and Polynesian ancestry inferred by SOURCEFIND.**
(A) Each individual is colored according to their island of residence.
(B) The black points indicate the individuals living in villages where Polynesian languages are spoken, the rest of the samples are colored in gray. The blue line indicates the regression line considering all the individuals from Vanuatu. The gray area indicates the 95% confidence level.

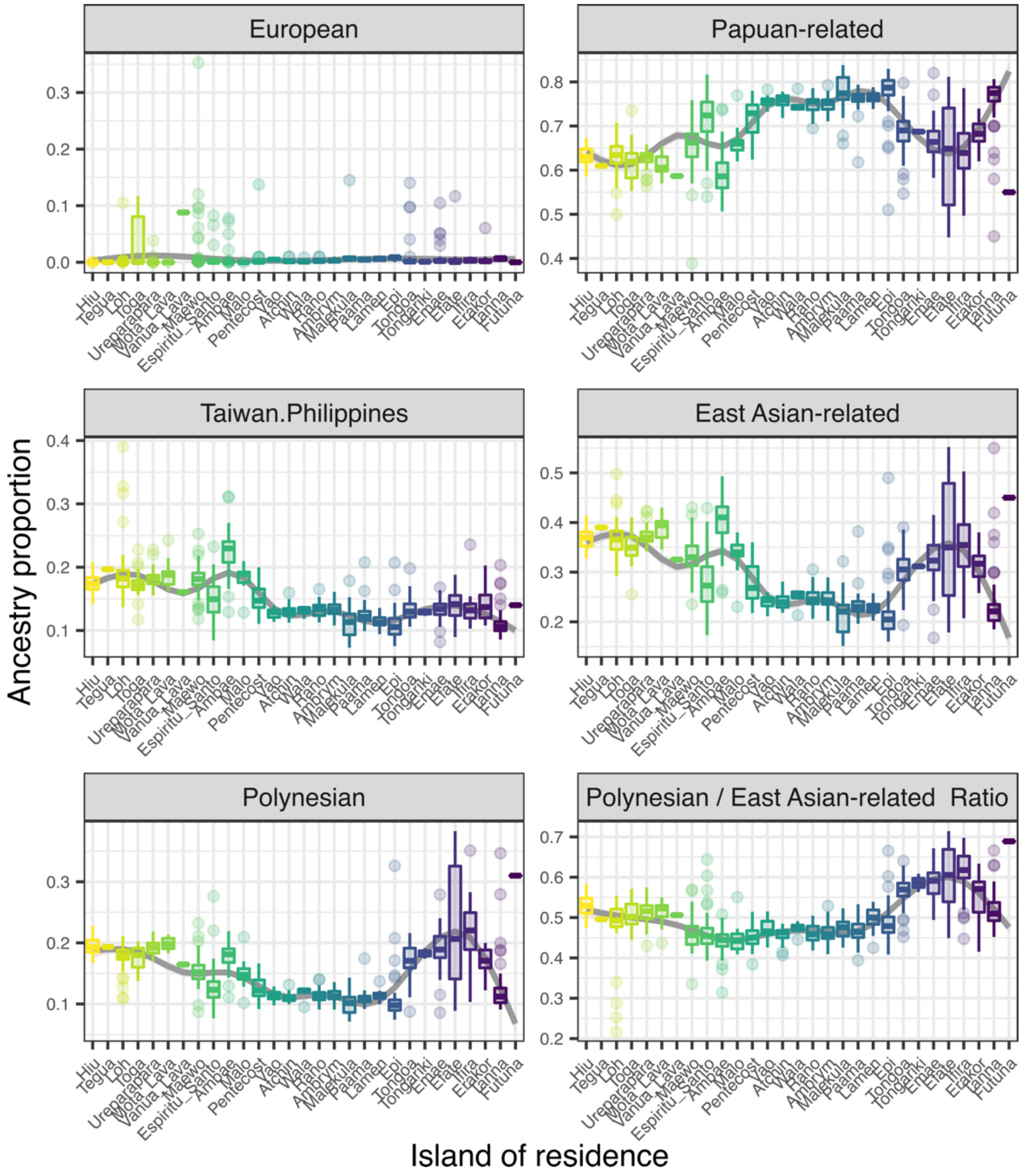


**Figure S10. Distributions of ancestry proportions in ni-Vanuatu, estimated by SOURCEFIND, for each island and ancestry.**

The line, box, whiskers and points, respectively, indicate the median, interquartile range (IQR), 1.5*IQR and outliers. Colors indicate the geographical latitude coordinates of each island, as in Fig. 1A.

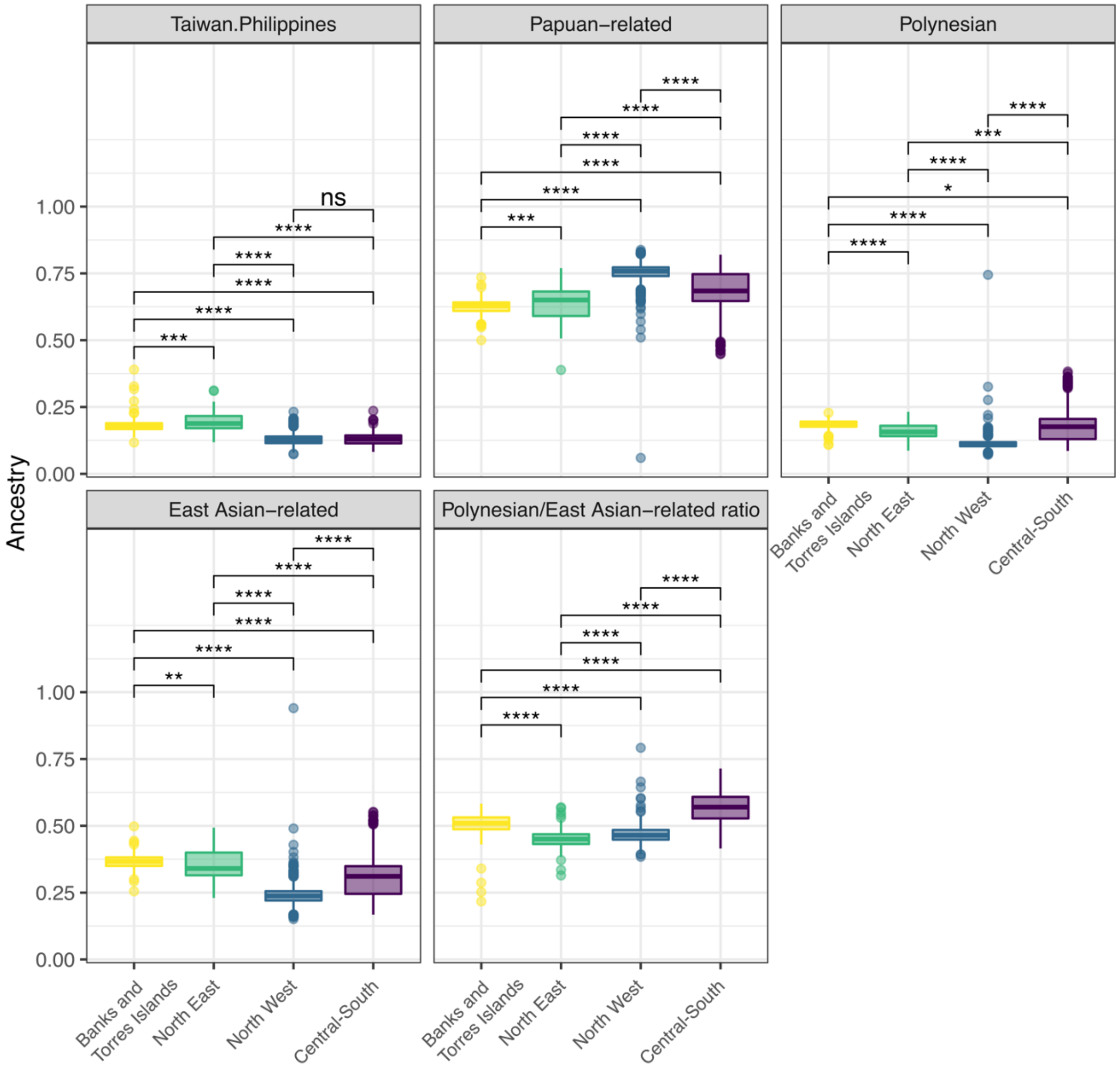


**Figure S11. Distributions of ancestry proportions of ni-Vanuatu among genetic clusters, according to fineSTRUCTURE at $K_{FS}$ = 4.**
The line, box and whiskers, respectively, indicate the median, interquartile range (IQR) and 1.5*IQR. We tested significant differences in ancestry between each cluster using a Wilcoxon test with Bonferroni correction. *$P < 0.05$, **$P < 0.01$, ***$P < 0.001$, ****$P < 0.0001$, ns=not significant.

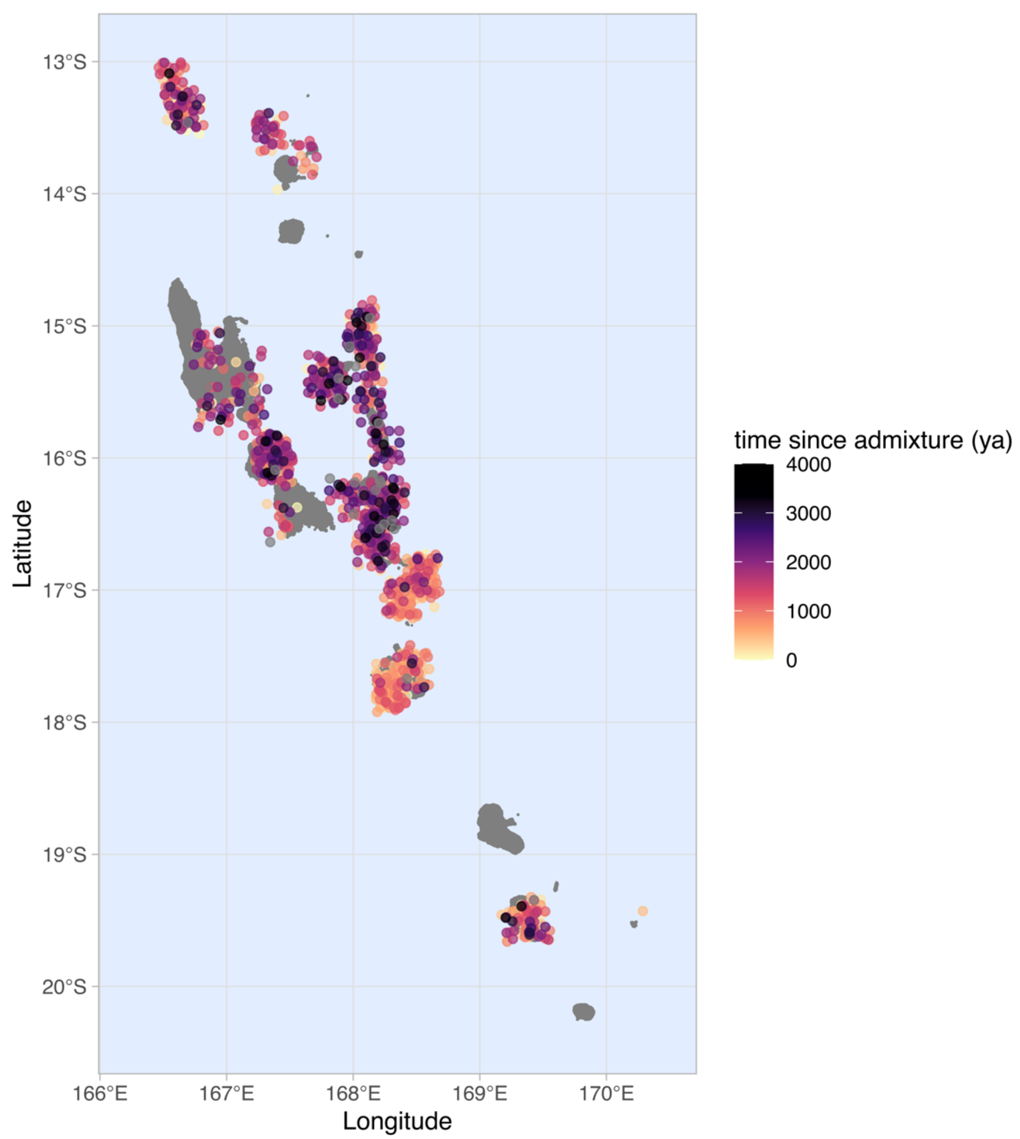

**Figure S12. Map showing admixture dates estimated per individual with GLOBETROTTER.**

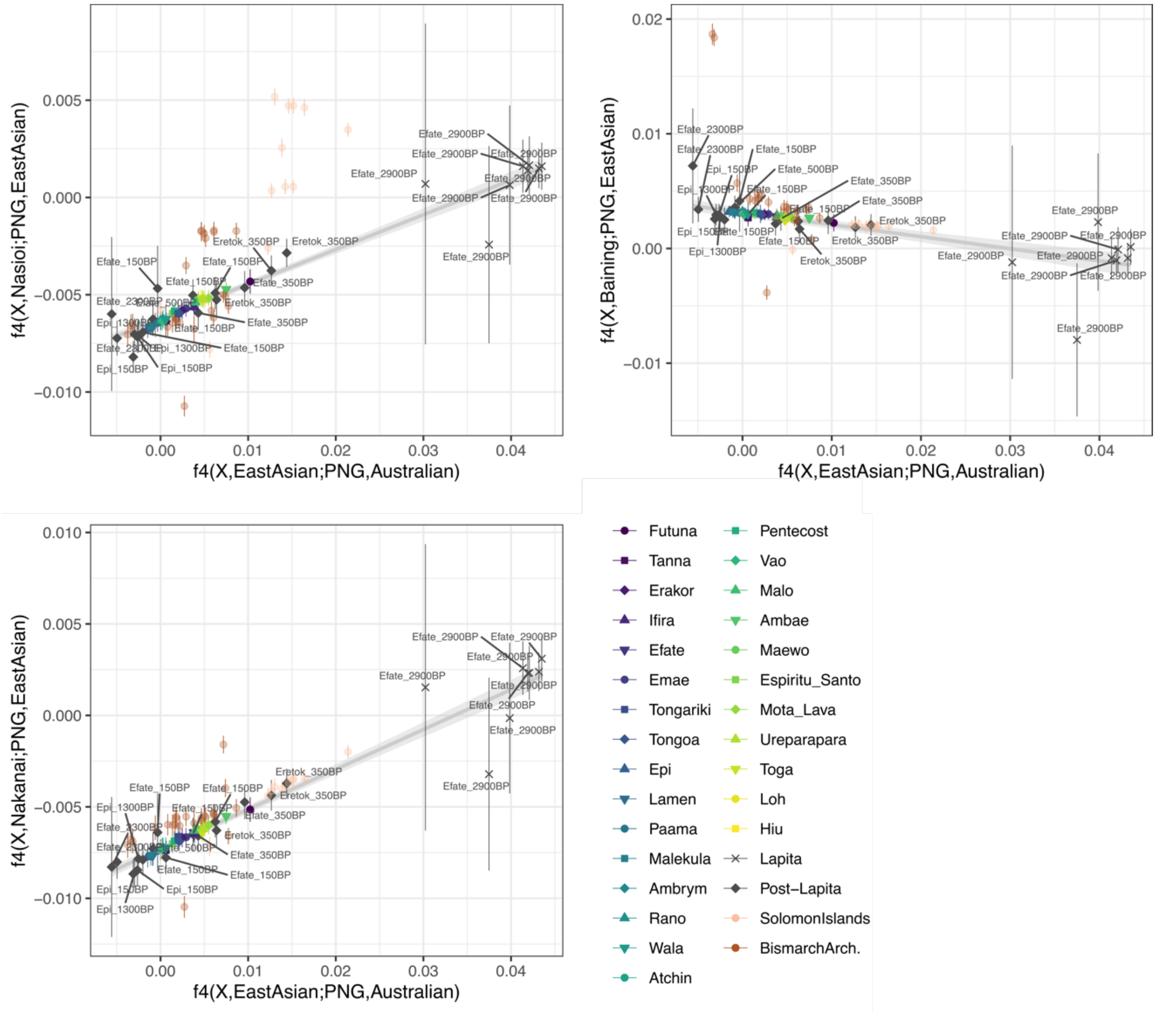


**Figure S13. Allele sharing of ni-Vanuatu with Near Oceanians, according to $f_4$-statistics.** The x-axis indicates $f_4$(X, New Guinean Highlanders; EastAsian, Australian), which measures allele sharing of ni-Vanuatu and Near Oceanians (X) with East Asians. The y-axis indicates $f_4$(X, New Guinean Highlanders; Near Oceanian, EastAsian), which measures allele sharing of ni-Vanuatu and Near Oceanians (X) to each Near Oceanian population tested (Nakanai, Nasioi, Baining), relative to their allele sharing with New Guinean Highlanders. The regression line is estimated considering only modern individuals from Vanuatu. The bars show two standard errors. Deviations upper from the regression line indicate a higher affinity with the Near Oceanian population tested.

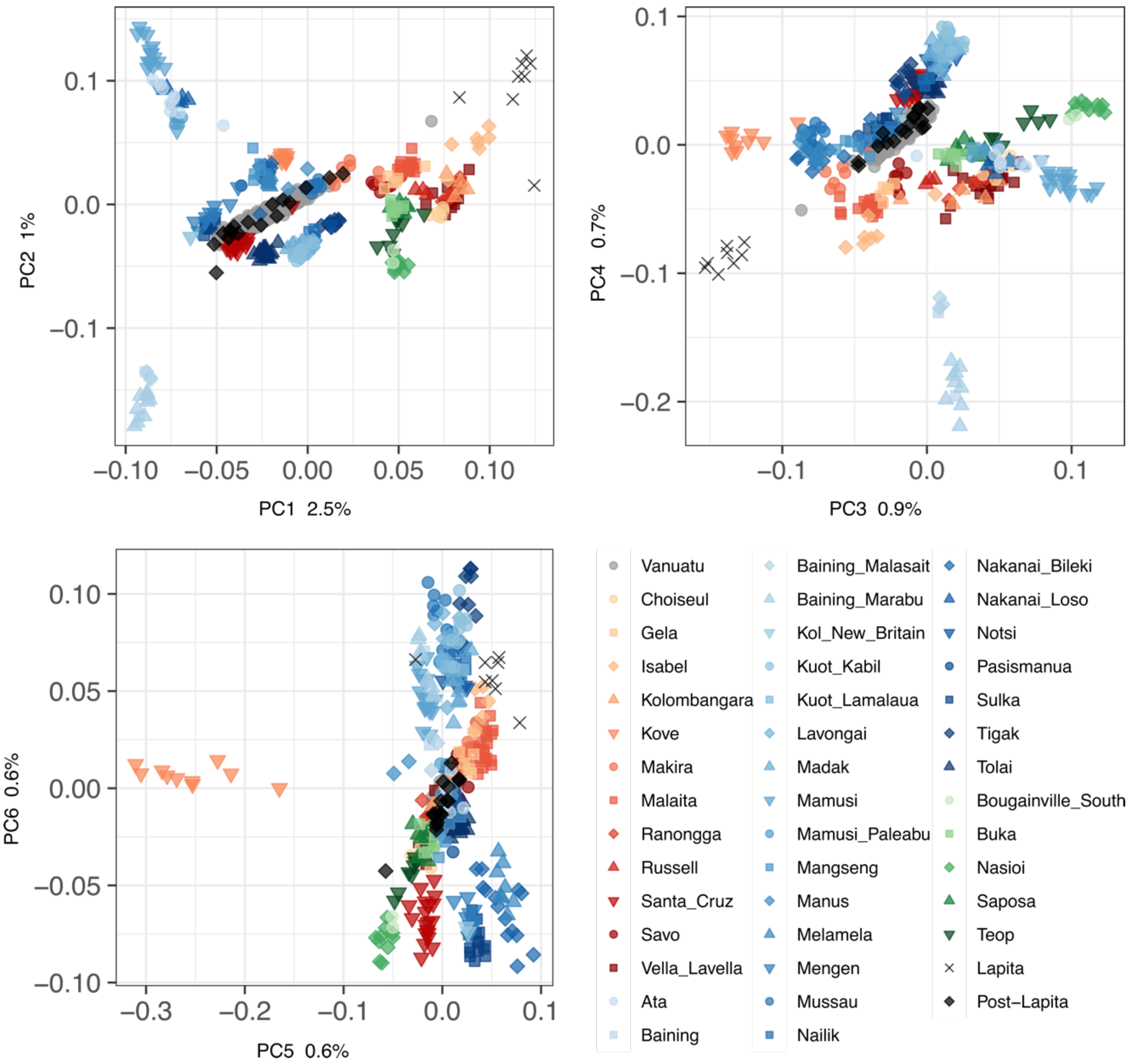


**Figure S14. Principal Component Analysis (PCA) of ni-Vanuatu, together with other populations from Near Oceania.**
The modern and ancient samples from Vanuatu were projected into a PCA of a previous SNP array data set [41].

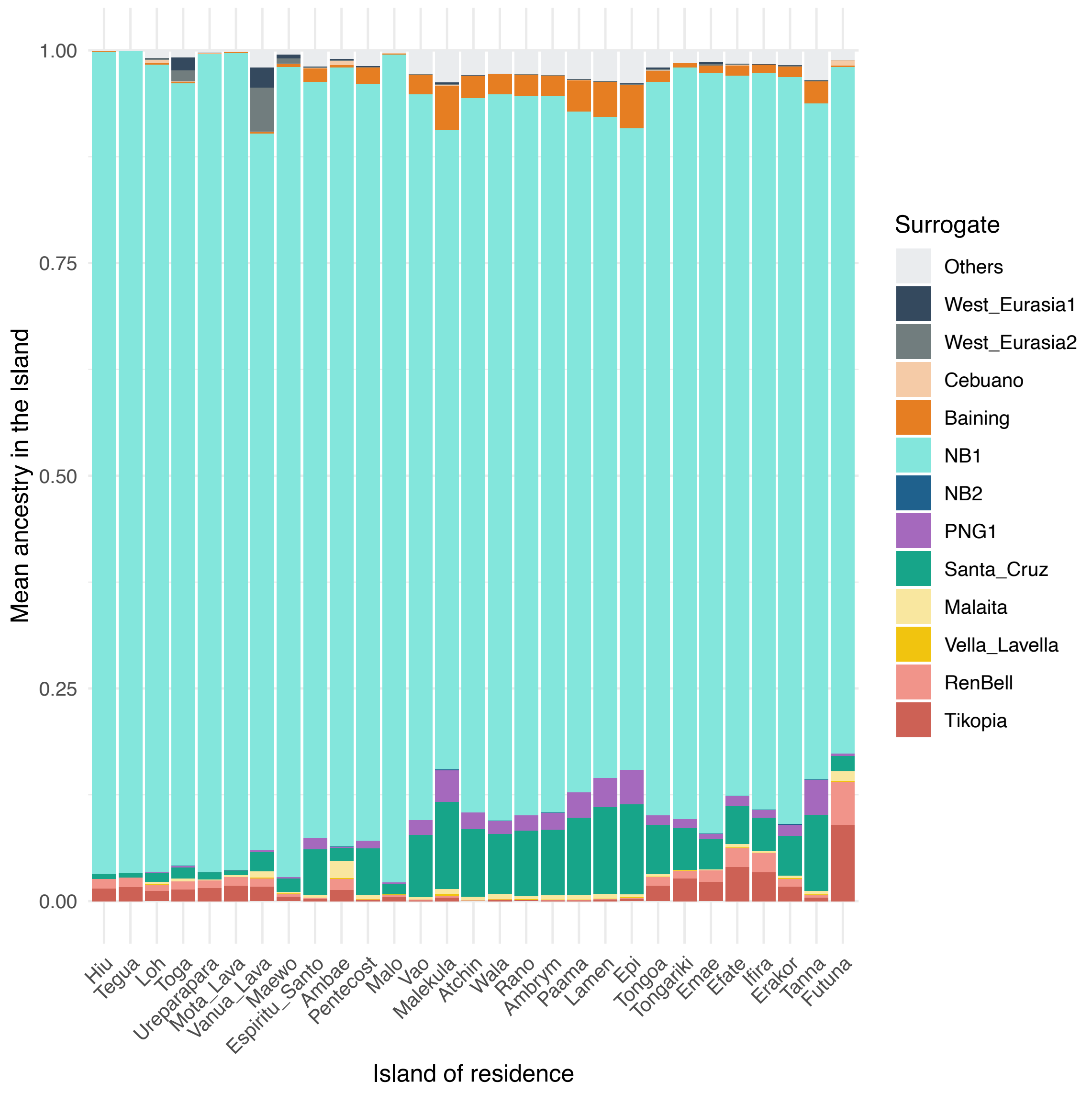


**Figure S15. Ancestry proportions in ni-Vanuatu, estimated by SOURCEFIND, considering all the reference populations as possible surrogates (Table S4).**
The bar plot indicates the mean value across individuals, for each ancestry in each island of residence.

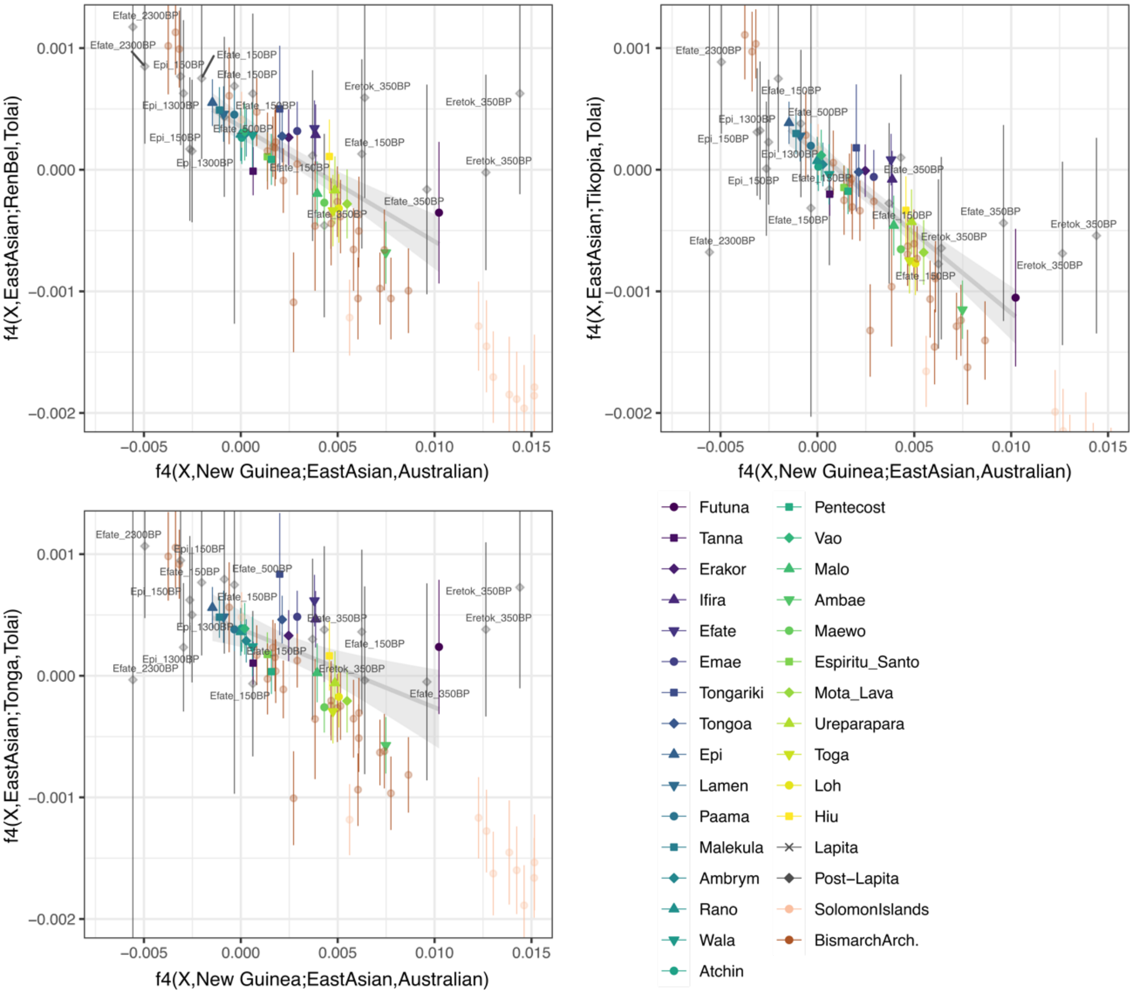


**Figure S16. Allele sharing of ni-Vanuatu with Polynesians, according to $f_4$-statistics.**
The x-axis indicates $f_4$(X, New Guinean Highlanders; EastAsia, Australian), which measures allele sharing of ni-Vanuatu and Near Oceanians (X) with East Asians. The y-axis indicates $f_4$(X, EastAsian; Polynesian, Tolai), which measures allele sharing of ni-Vanuatu and Near Oceanians (X) with Polynesians (Rennell and Bellona [RenBel], Tikopia, Tonga), relative to their East Asian ancestry. The regression line is estimated considering only modern and ancient individuals from Vanuatu. The bars show two standard errors. Deviations upper from the regression line indicate a higher affinity with the Polynesian population tested.

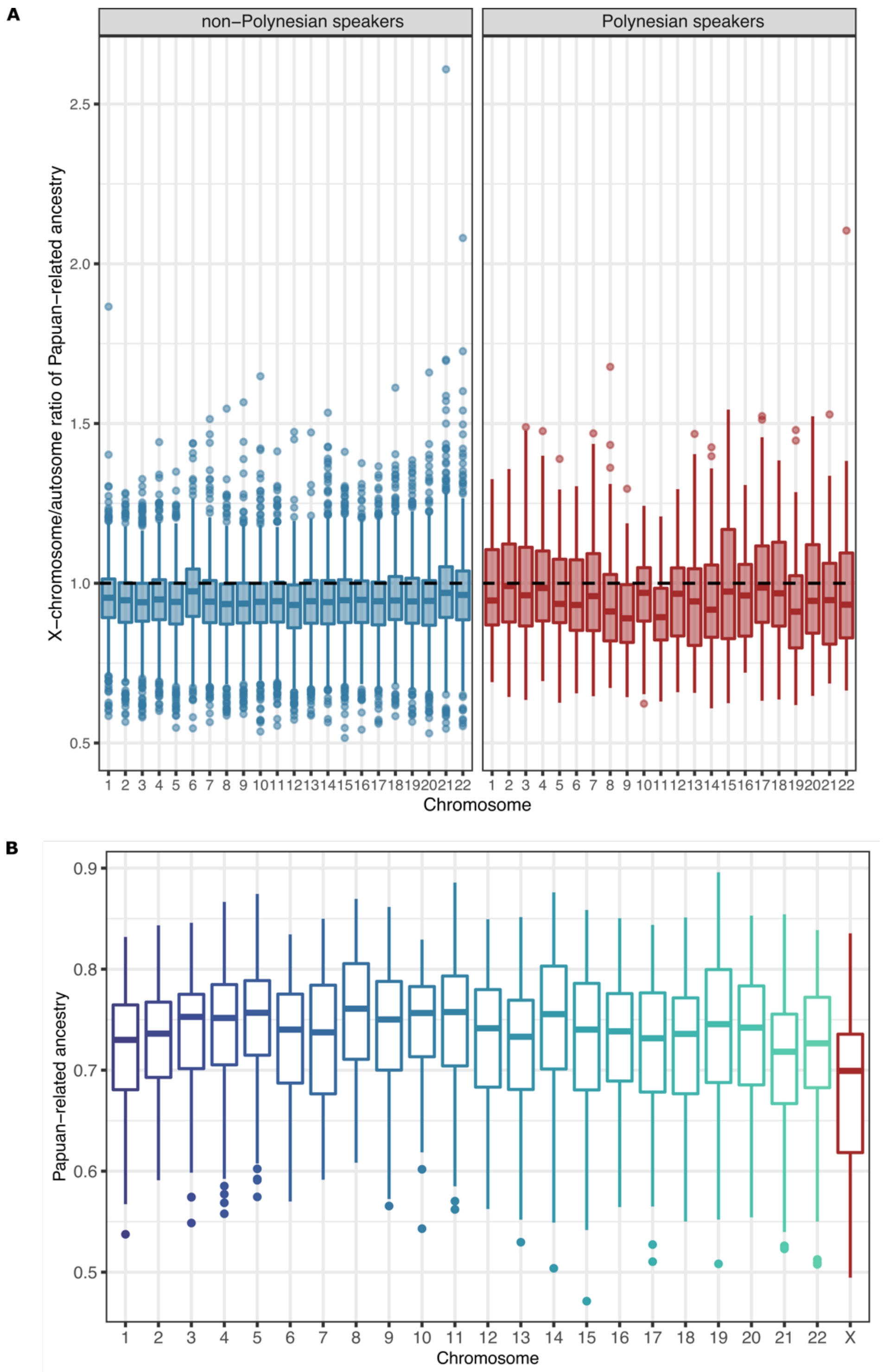


**Figure S17. Sex-biased admixture in Vanuatu.**
(A) X-to-autosome ratio of Papuan-related ancestry proportions in ni-Vanuatu who speak non-Polynesian or Polynesian languages, estimated using the SNP array data (Wilcoxon test *P*-value $< 1.36\times10^{-5}$, after Bonferroni correction).
(B) Papuan-related ancestry proportions in ni-Vanuatu, estimated for the 22 autosomes and the X chromosome separately, using the high-coverage genome sequencing data obtained for a subset of 179 ni-Vanuatu [18].

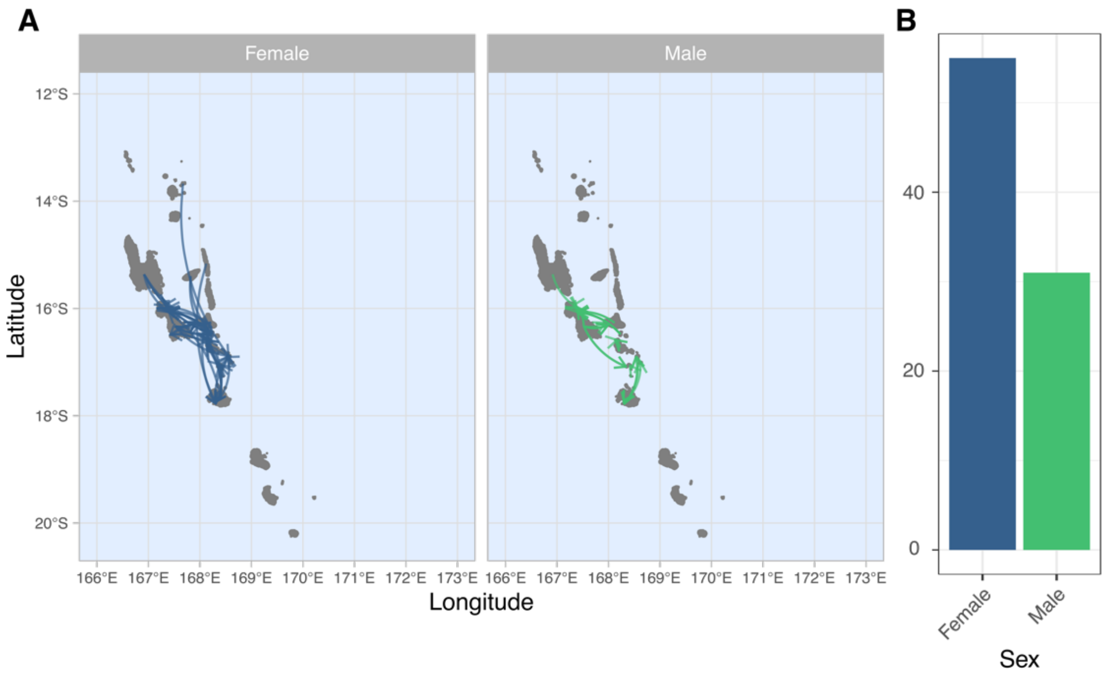


**Figure S18. Self-reported migrations among ni-Vanuatu spouses.**
(A) Map of self-reported migrations for females (blue) and males (green), separately. The arrows connect the place of birth of individuals to their place of residence.
(B) Barplot showing the number of females (blue) and males (green) among ni-Vanuatu spouses who reported to migrate during their lifetime.

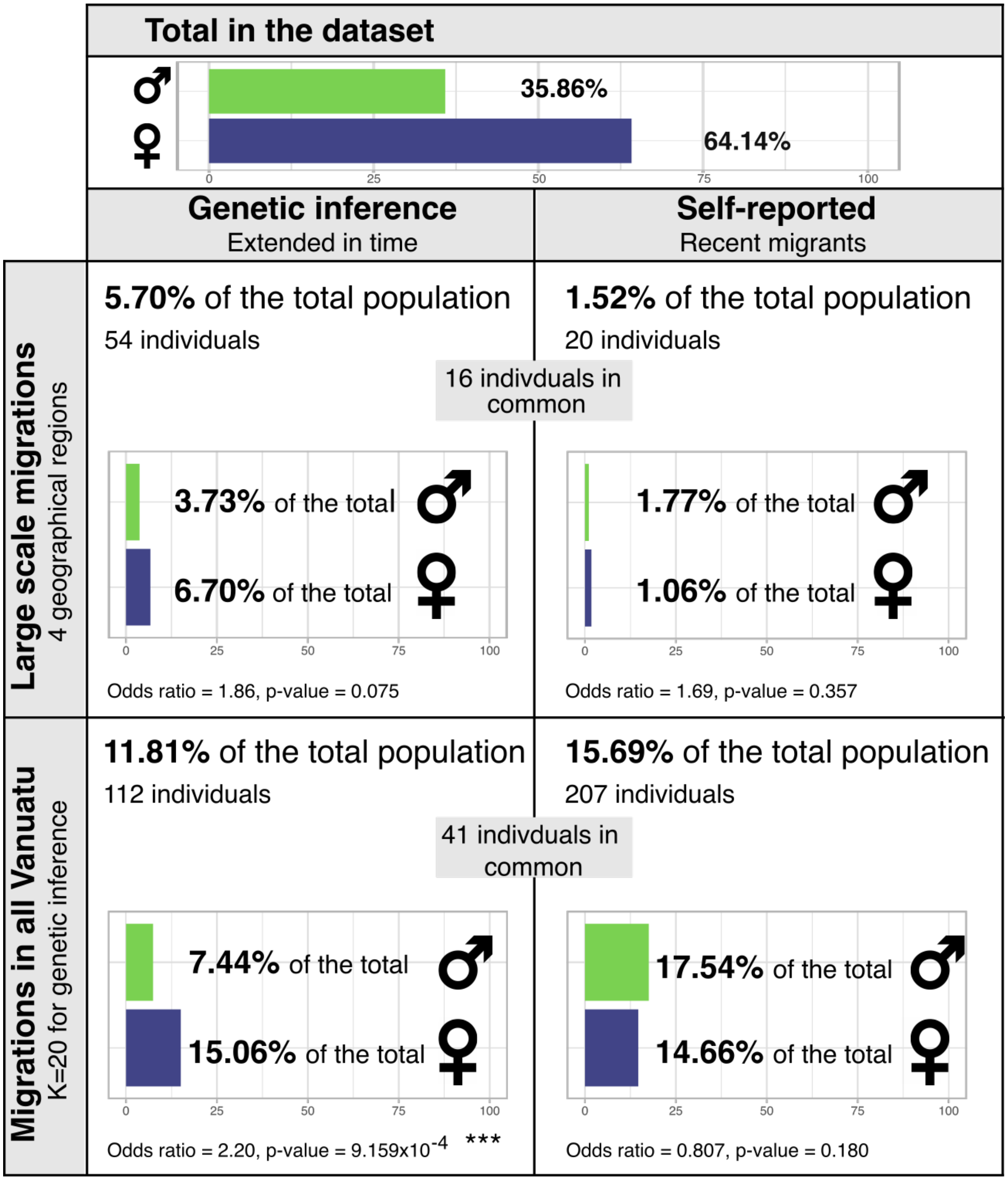


**Figure S19. Proportions of female and male ni-Vanuatu migrants, based on genetic clusters or self-reported information.**

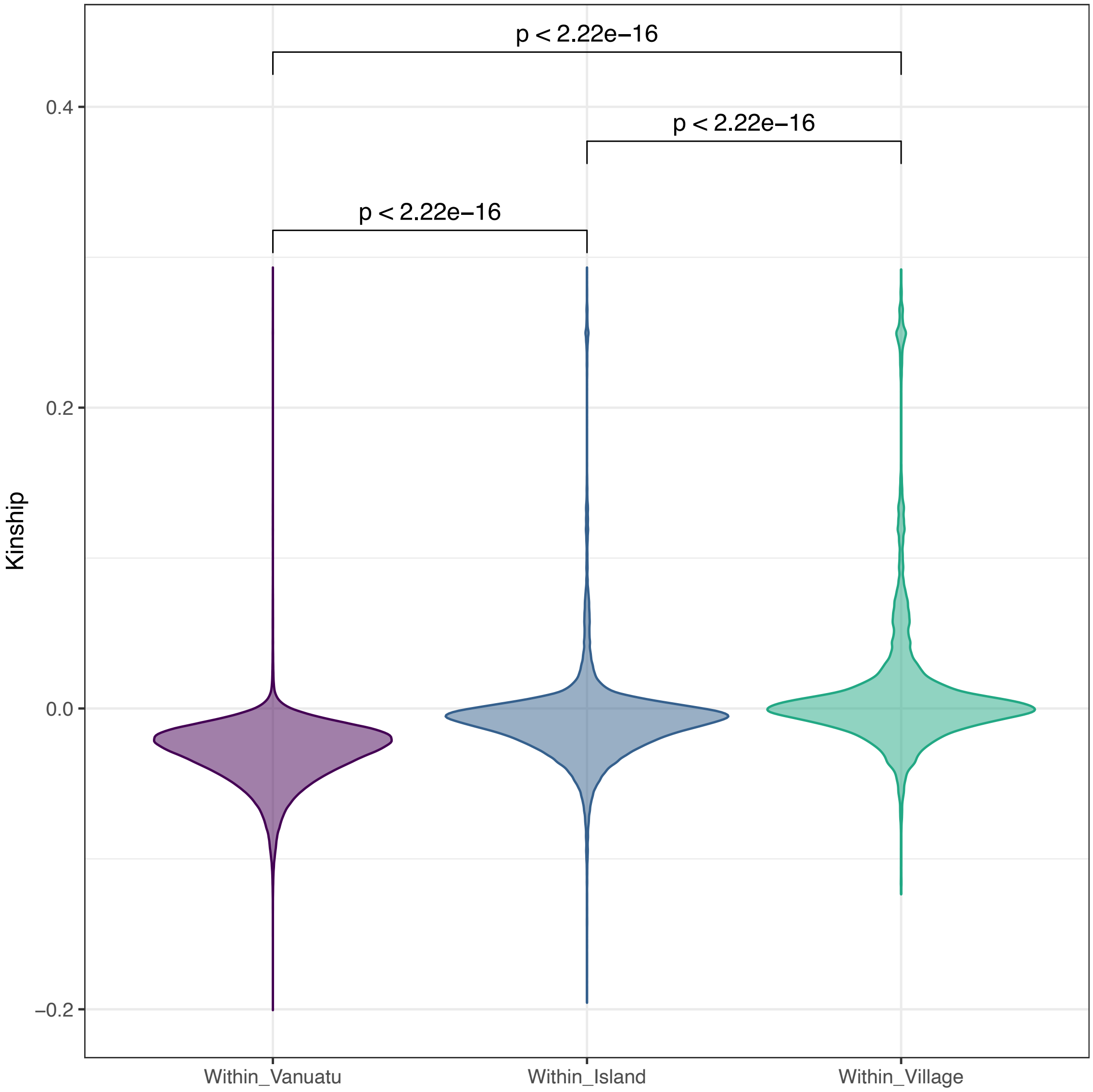


**Figure S20. Comparison of kinship levels among ni-Vanuatu living in the same village, the same island or the entire archipelago.**
Significance of the differences was tested by a Wilcoxon test.

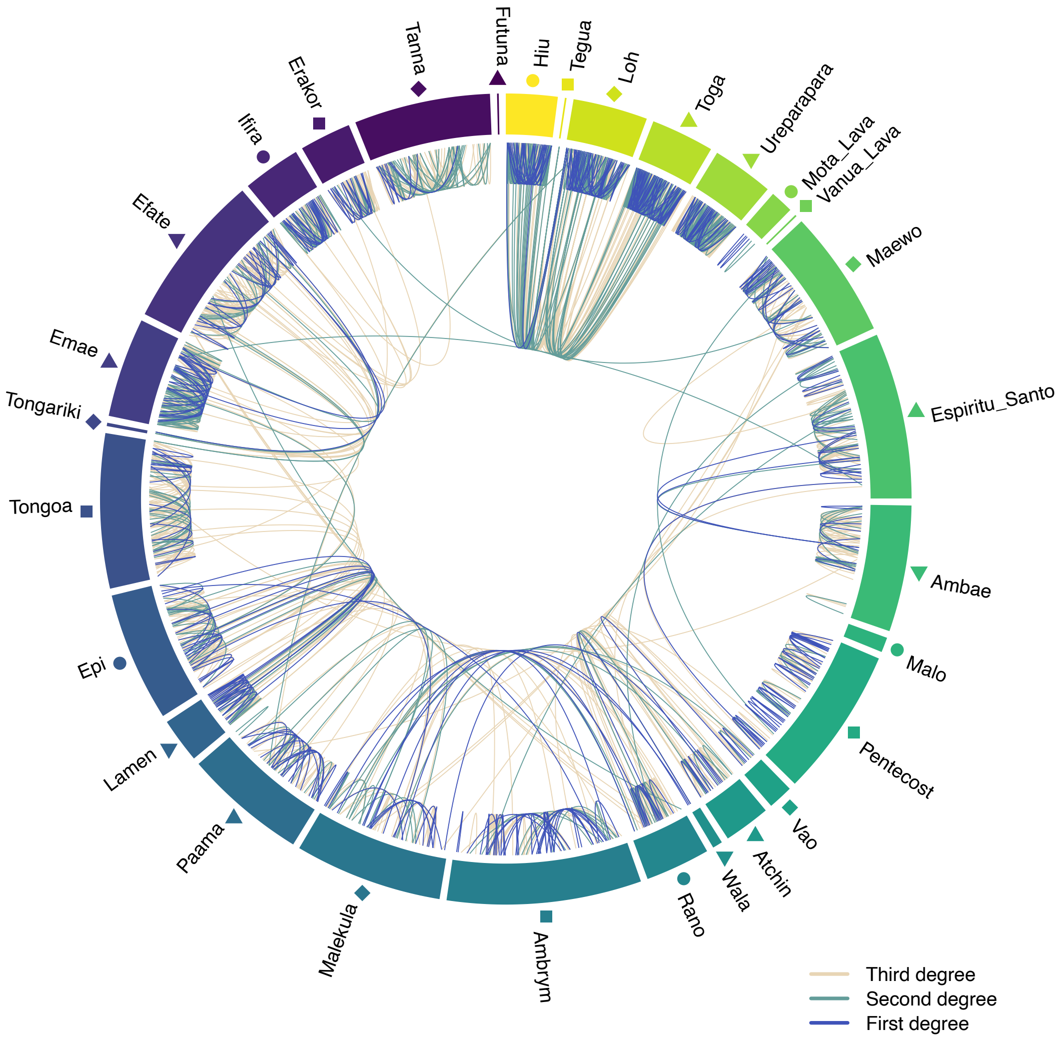


**Figure S21. Genetic relatedness and islands of residence of related ni-Vanuatu.**
Each line connects two individuals who are related up to third degree. Islands are order from north to south (clockwise).

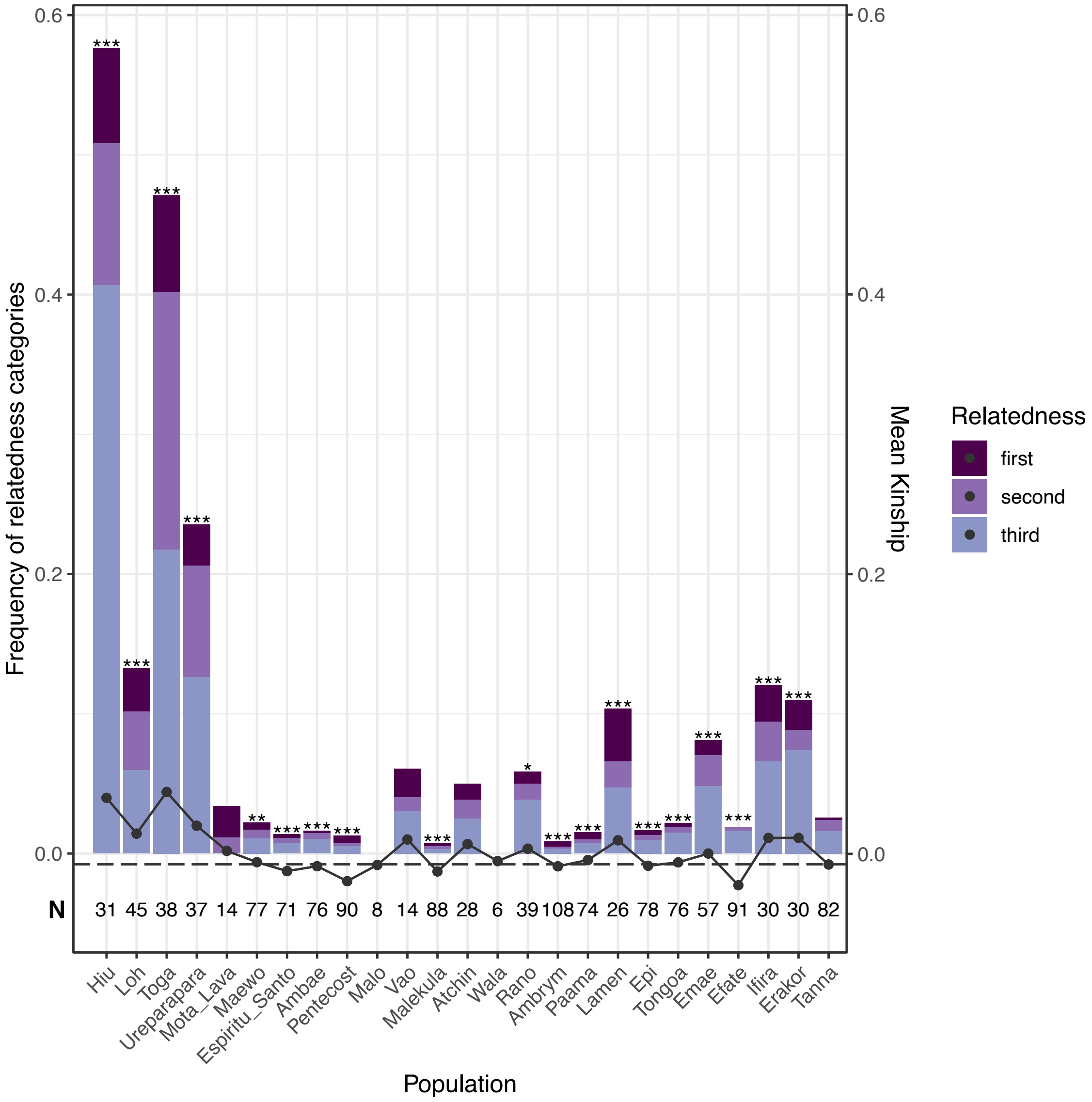


**Figure S22. Degrees of genetic relatedness of ni-Vanuatu from the different Vanuatu islands.** Bar plots indicate the proportion of pairs of related individuals according to the inferred degree of genetic relatedness, in each island. The black points and indicate the mean kinship per island. Significance was tested by a Fisher's exact test. **P* < 0.05, ***P* < 0.01, ****P* < 0.001.

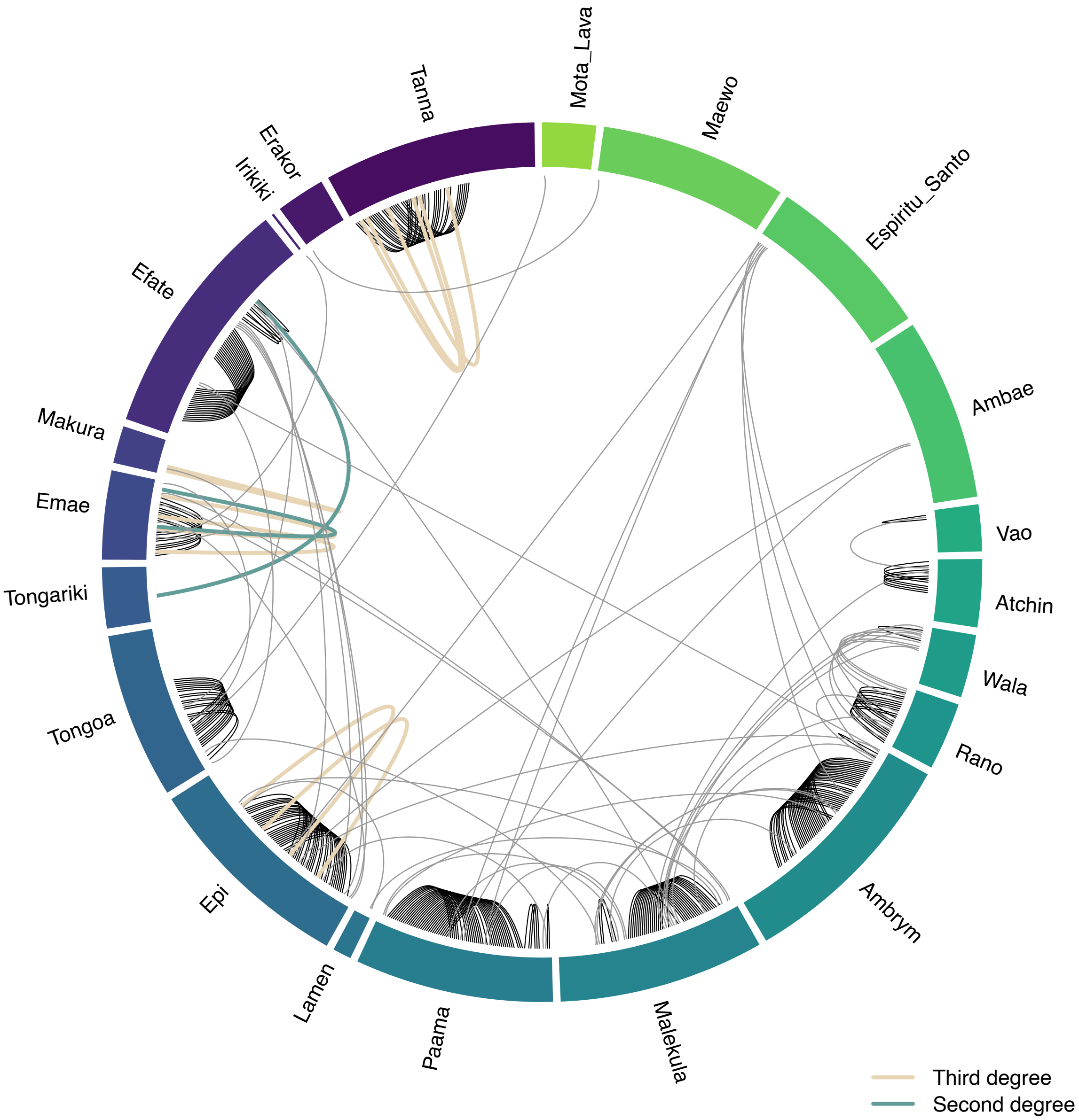


**Figure S23. Genetic relatedness and islands of residence of ni-Vanuatu spouses.**
Each line connects the two spouses, and the color indicates the degree of genetic relatedness. Islands are order from north to south (clockwise).

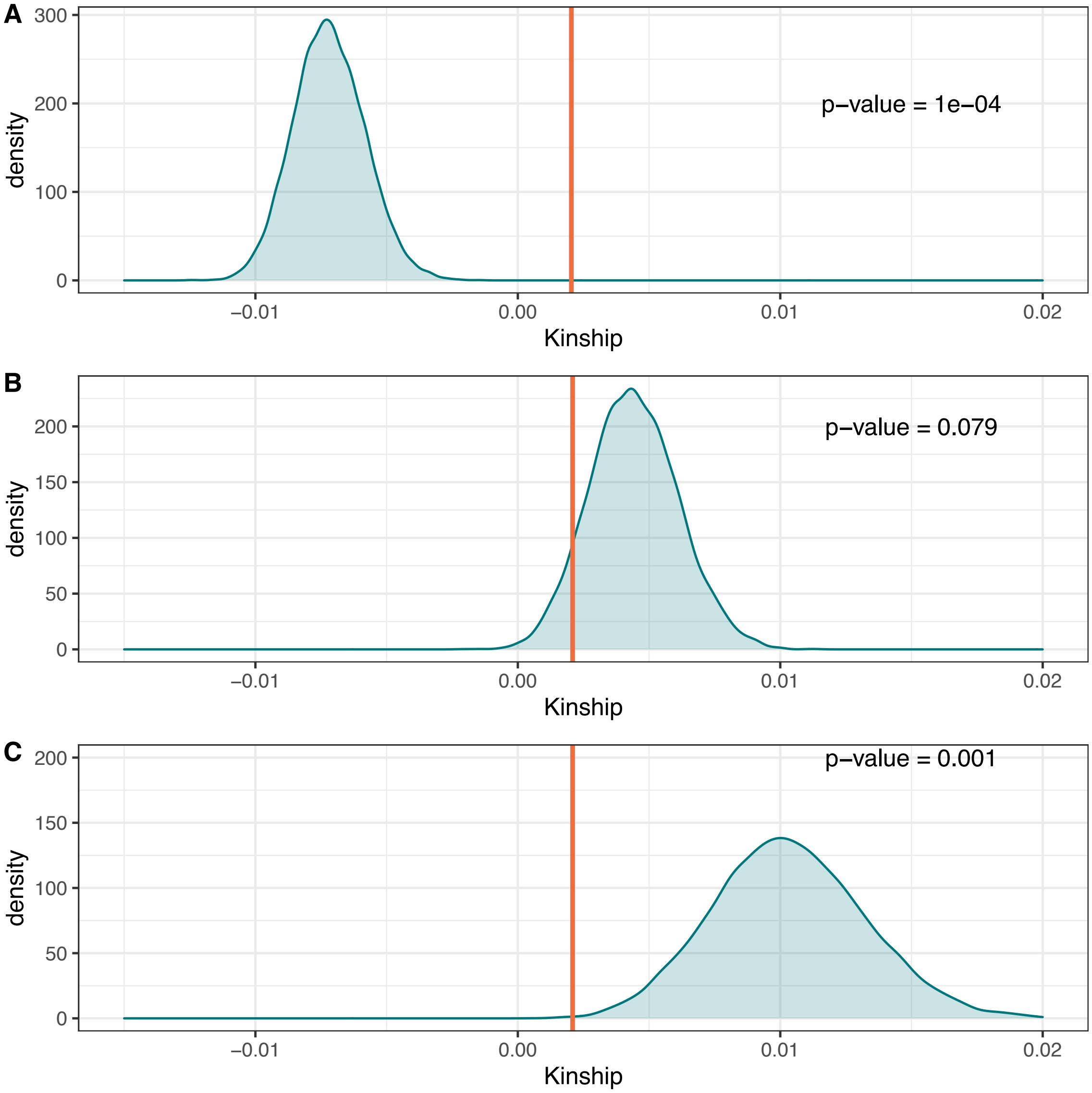


**Figure S24**. **Kinship among spouses and random pairs of individuals.**
Kinship coefficients among spouses and random pairs of individuals from the same (A) island (excluding first-degree related pairs), (B) village (excluding first-degree related pairs) and (C) village (including first-degree related pairs). The orange vertical line indicates the average kinship coefficient among spouses. The blue density curve indicates the null distribution of kinship coefficients estimated from the sampling of random pairs of individuals. Each data point of the null distribution is estimated as the mean kinship coefficient in randomly sampled pairs of individuals. Of note, negative kinship coefficients are expected when the two compared individuals are from different populations.

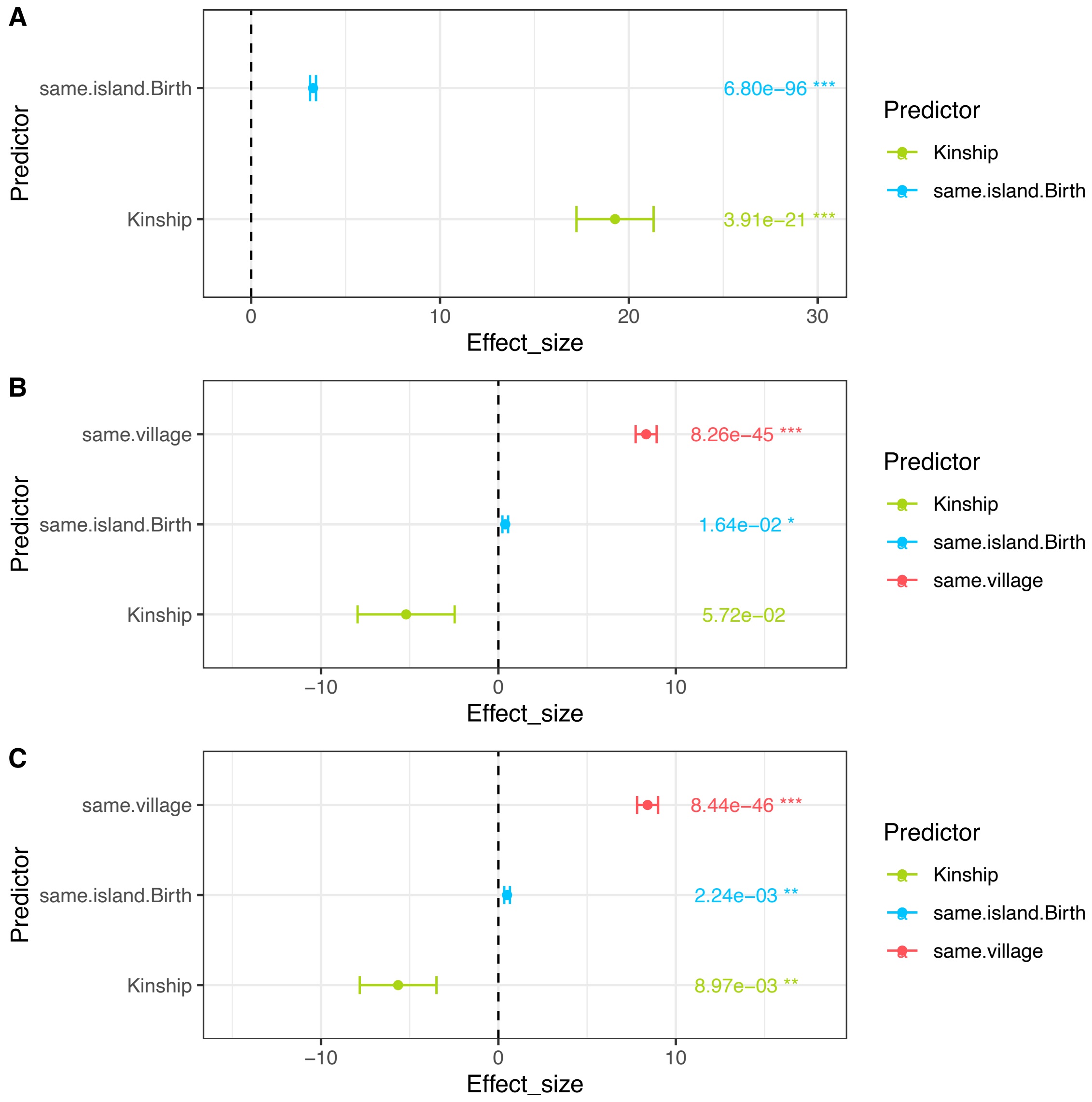


**Figure S25. Effects of geographical location and kinship on mate choice.**
(A) Effect sizes estimated when not including the village of residence as a predictor.
(B) Effect sizes estimated when including the village of residence as a predictor and excluding first-degree related pairs from the analysis.
(C) Effect sizes estimated when including the village of residence as a predictor and including first-degree related pairs from the analysis.
(A-C) Effect sizes of the logistic regression modelling the probability of mating as a function of three predictors: the island of birth, the village of residence and kinship coefficients.

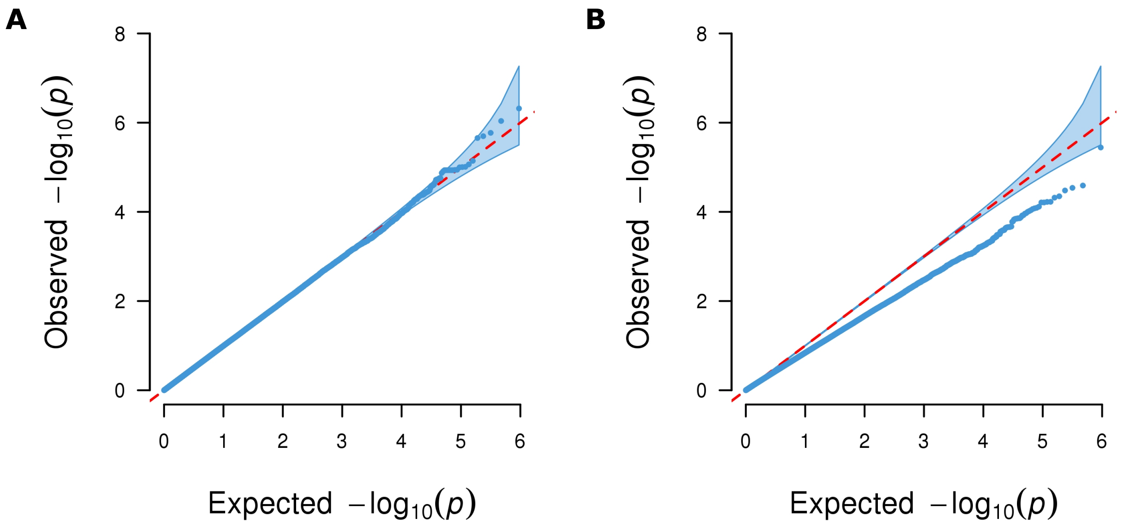


**Figure S26. Q-Q plots of observed and expected *P*-values of a regression model testing for SNP-based assortative mating.**

(A) *P*-values are from a model that does not control for population structure.

(B) *P*-values are from a model that controls for population structure (island of birth, kinship and ancestry). The shaded area indicates the 95% confidence interval.
